# SARS-CoV-2 infection in the Syrian hamster model causes inflammation as well as type I interferon dysregulation in both respiratory and non-respiratory tissues including the heart and kidney

**DOI:** 10.1101/2021.04.07.438843

**Authors:** Magen E. Francis, Una Goncin, Andrea Kroeker, Cynthia Swan, Robyn Ralph, Yao Lu, Athema L. Etzioni, Darryl Falzarano, Volker Gerdts, Steven Machtaler, Jason Kindrachuk, Alyson A. Kelvin

**Affiliations:** Vaccine and Infectious Disease Organization VIDO, University of Saskatchewan, Saskatoon, SK, S7N 5E3, Canada; Department of Biochemistry, Microbiology, and Immunology, University of Saskatchewan, Saskatoon, SK, S7N 5E5, Canada; Department of Medical Imaging, University of Saskatchewan, Saskatoon, SK, S7N 0W8, Canada; Department of Veterinary Microbiology, Western College of Veterinary Medicine, University of Saskatchewan, Saskatoon, SK, S7N 5B4, Canada; Department of Pathobiology, Tuskegee University College of Veterinary Medicine, 1200 W. Montgomery Road, Tuskegee Institute, Alabama, 36088, USA; Laboratory of Emerging and Re-Emerging Viruses, Department of Medical Microbiology, University of Manitoba, Winnipeg, MB, R3E 0J9, Canada; Department of Microbiology and Immunology, Faculty of Medicine, Dalhousie University, Halifax, NS B3H 4R2, Canada; Department of Pediatrics, Division of Infectious Disease, Faculty of Medicine, Dalhousie University, Halifax, NS B3K 6R8, Canada; Canadian Centre for Vaccinology, IWK Health Centre, Halifax, NS B3K 6R8, Canada

## Abstract

COVID-19 (coronavirus disease 2019) caused SARS-CoV-2 (severe acute respiratory syndrome coronavirus 2) infection is a disease affecting several organ systems. A model that captures all clinical symptoms of COVID-19 as well as long-haulers disease is needed. We investigated the host responses associated with infection in several major organ systems including the respiratory tract, the heart, and the kidneys after SARS-CoV-2 infection in Syrian hamsters. We found significant increases in inflammatory cytokines (IL-6, IL-1beta, and TNF) and type II interferons whereas type I interferons were inhibited. Examination of extrapulmonary tissue indicated inflammation in the kidney, liver, and heart which also lacked type I interferon upregulation. Histologically, the heart had evidence of mycarditis and microthrombi while the kidney had tubular inflammation. These results give insight into the multiorgan disease experienced by people with COVID-19 and possibly the prolonged disease in people with post-acute sequelae of SARS-CoV-2 (PASC).

## Introduction

The SARS-CoV-2 (severe acute respiratory coronavirus 2) which causes COVID-19 (coronavirus 2019) emerged in late 2019 with subsequent massive public health and economic impacts^1,2^. SARS-CoV-2 is a positive-sense, single-stranded RNA virus of family *Coronoviridae*, genus *Betacoronavirus*, and since its emergence has led to 124 million confirmed cases and 2.7 million deaths worldwide^3,4^. Although coronavirus infections in humans were previously thought to only cause mild, cold-like symptoms, the more recent coronaviruses to emerge, SARS-CoV-1 and MERS-CoV (Middle East Respiratory Syndrome coronavirus), are associated with a wide array of respiratory and non-respiratory symptoms^5,6^, ranging from mild to severe, leading to organ failure and death^7^. Asymptomatic cases have also been reported^8^. Given the burden on public health, as well as the range of disease severity, developing animal models in order to study SARS-CoV-2 pathogenesis and identify countermeasures is of high importance. *In vivo* assessment of host response to viral infection is essential to identify host factors regulating severe disease as well as biomarkers that may help inform patient supportive care measures.

Assessment of host response to SARS-CoV-2 has led to valuable insight into disease progression and possible immune mechanisms leading to severe disease. It has now been widely shown that severe cases of COVID-19 in humans are often accompanied by increased expression of inflammatory genes, including Interleukin-6 (IL-6), Interleukin-1 beta (IL-1beta) and Tumour Necrosis Factor (TNF)^11–13^. Prior to SARS-CoV-2, it was known that coronaviruses were capable of regulating the type I interferon response^14,15^. Several studies have found that type I interferon responses are significantly blunted during SARS-CoV-2 infection suggesting this may also contribute to poor COVID-19 outcomes^16,17^. These clinical data suggest potential markers of pathogenesis for further experimental investigations.

Syrian hamsters were one of a handful of animal species identified early during the pandemic to be productively infected with SARS-CoV-2 and of potential use for preclinical development and *in vivo* analysis^18^. Hamsters were originally determined to be susceptible to SARS-CoV-1^19^ and since have been shown to be an ideal model for SARS-CoV-2^20^. Hamsters are able to support SARS-CoV-2 replication within the lower respiratory tract ^20–22^ with similar infection and disease dynamics to humans^23^, in contrast to ferrets and mice^24^. Further, hamsters mount an effective antibody response and have been utilized for both COVID-19 vaccine and treatment studies^21,22,25^. As with other non-traditional animal models, there is a lack of hamster-specific biological reagents for characterizing immune responses and the mechanisms of pathogenesis^26^.

Here, we investigated the immune response and pathogenesis of SARS-CoV-2 infection in hamsters to determine disease mechanisms and the involvement of specific organs such as the lungs, heart, and kidney since COVID-19 is a multisystem disease. SARS-CoV-2 inoculated Syrian hamsters were sampled over a 15-day time course for virological and immunological analysis by qRT-PCR, transcriptome analysis, virus titration assays, and immunohistochemistry (IHC). The respiratory tract had a high burden of replicating virus in both the upper and lower regions with specific blunting of the type I interferon response. While we found little evidence of infectious virus outside the respiratory tract, viral RNA was present in several organs including the kidneys, heart, lymph node, and liver with concomitant expression of inflammatory mediators such as IL-6, TNF, and IL-1beta. The lung, as well as extrapulmonary organs, showed signs of inflammatory cell infiltration leading to tissue damage. Taken together, we have defined the hamster model of immunopathogenesis in the context of COVID-19 which can be used to identify correlates of protection and disease, to evaluate evaluation of medical countermeasures, as well as to dive into the mechanisms of COVID-19 and related dysfunction such as post-acute sequelae of SARS-CoV-2 (PASC), also known as long COVID^27^.

## Results

### SARS-CoV-2-inoculated hamsters have weight loss that correlated with viral load, infectious virus in the respiratory tract, and viral RNA in non-respiratory tissues

Syrian hamsters are susceptible to infection with SARS-CoV-2 suggesting they may be valuable for the investigation of immunopathology and mechanisms of disease^22^. Hamsters aged 6-8 weeks were inoculated intranasally with SARS-CoV-2 at 10^5^ TCID50. The animals were monitored for temperature and weight throughout the course of the infection. As typical during viral infection in small animals, hamsters began showing a decrease in temperature by day 1 post inoculation (pi) (**Figure 1A**) which returned to baseline between day 3 and 7 (**Figure 1B**). Inoculated hamsters began losing weight on day 2 pi, with weights being significantly different from baseline on days 3 to 7. Weight loss was most severe on day 6 pi where the average body weight of the inoculated animals was 88.26% of original weight (**Figure 1C**). After day 6, there was a steady increase in weight over the remaining time course on average recovering to baseline by day 12. Individual animals showed two different trajectories of weight loss: mild (∼4% weight loss) and severe (>15% weight loss) (**Figure 1D**). No animals lost ≥20% of original weight or reached endpoint criteria during the study.

**Figure 1.**
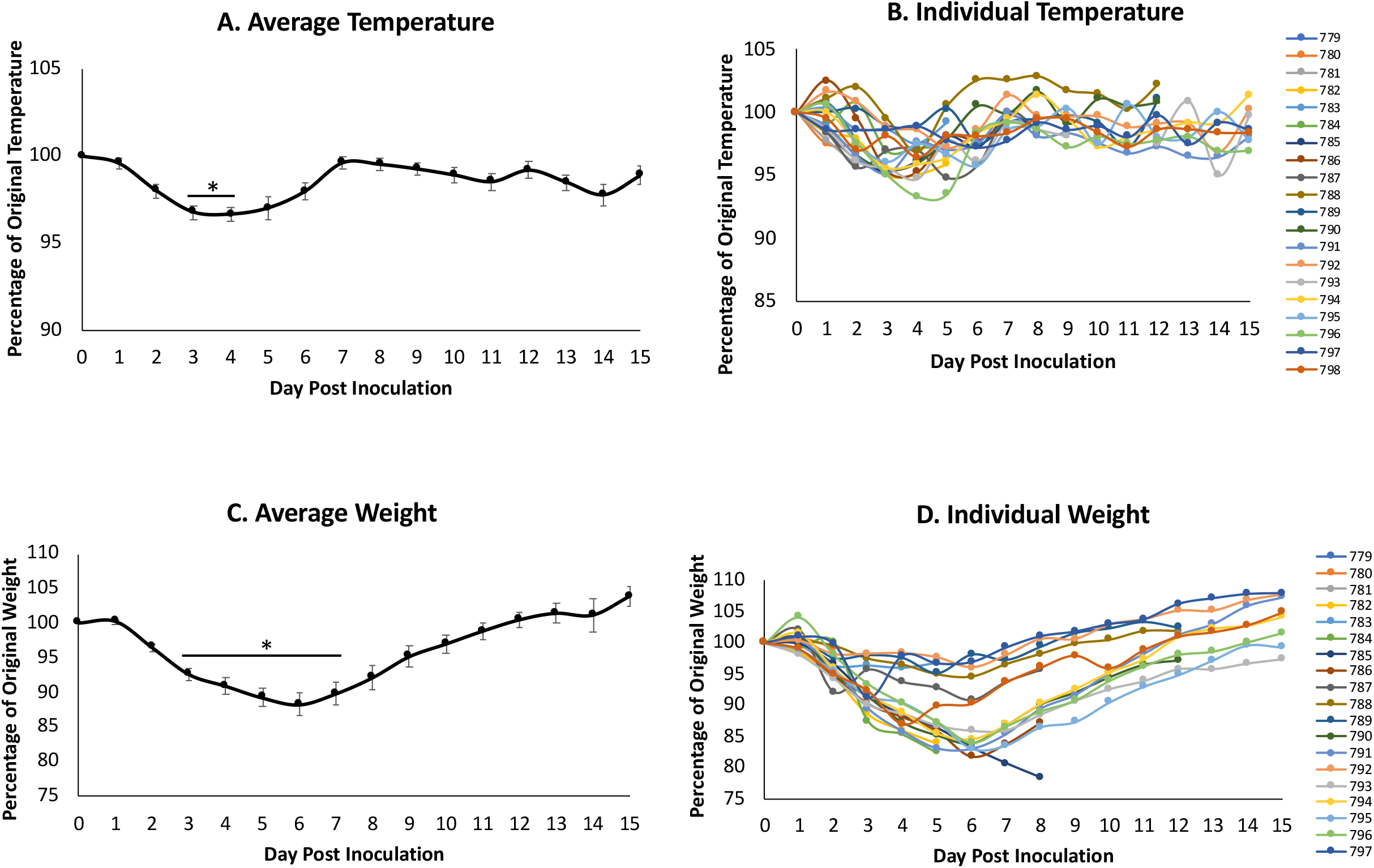
Intranasal SARS-CoV-2 infection in hamsters leads to significant temperature decrease and weight loss. Syrian hamsters were intranasally inoculated with the Severe Acute Respiratory Syndrome coronavirus 2 (SARS-CoV-2) at 10^5^ TCID_50_. Temperature (A and B) and weight (C and D) were recorded for 15 days post inoculation. Average results (A and C) show the mean. Individual results represent each animal (B and D). * indicates a p-value <0.05 determined by ANOVA comparing hamsters on the days post inoculation to baseline (day 0). Error bars indicate +/- standard error (SE).

COVID-19 affects several organ systems in humans including the heart, kidneys, liver, spleen and large intestine^31,32^. Therefore, we were interested in determining the tropism of the virus and associated immunopathology throughout the body of infected hamsters to gain insight into disease. Animals were removed from the study on days 0, 2, 5, 8, 12 and 15 pi to assess tissue-specific responses for necropsy and analysis of the immunopathology. Blood, nasal washes and tissues (respiratory and non-respiratory) were collected. Viral tropism was assessed by qRT-PCR for the SARS-CoV-2 envelope gene (E) and virus titration with TCID50 assays (**Figure 2**). E gene expression analysis is represented by brown circles on the graphs. Analysis of the respiratory tract revealed vRNA present in the nasal turbinates, trachea, and all four tested lung lobes (right cranial, right middle, right caudal, and accessory) on days 2, 5 and 8 pi. Viral RNA was still detected in the right middle lung lobe and the accessory lung lobe of one animal on day 12 pi. Live viral titer analysis, represented by the grey circles, indicated that infectious virus was present in the nasal turbinates on days 2 (7.5 TCID50/mL (Log10)) and 5 (6 TCID50/mL (Log10)) pi. Infectious virus peaked in the lungs on day 2, at ∼9 TCID50/mL (Log10) in the right middle lung and was present until day 5. The trachea had the lowest virus levels of all the respiratory tissue which was only present on day 2 (∼3 TCID50/mL (Log10)). vRNA was also detected in non-respiratory tissues including the heart, kidney, large intestine, and the mediastinal lymph node at low to moderate levels. The mediastinal lymph node had the highest level of vRNA, ∼5 TCID50 (Log10) and was the only non-respiratory tissue to be positive for infectious virus.

**Figure 2.**
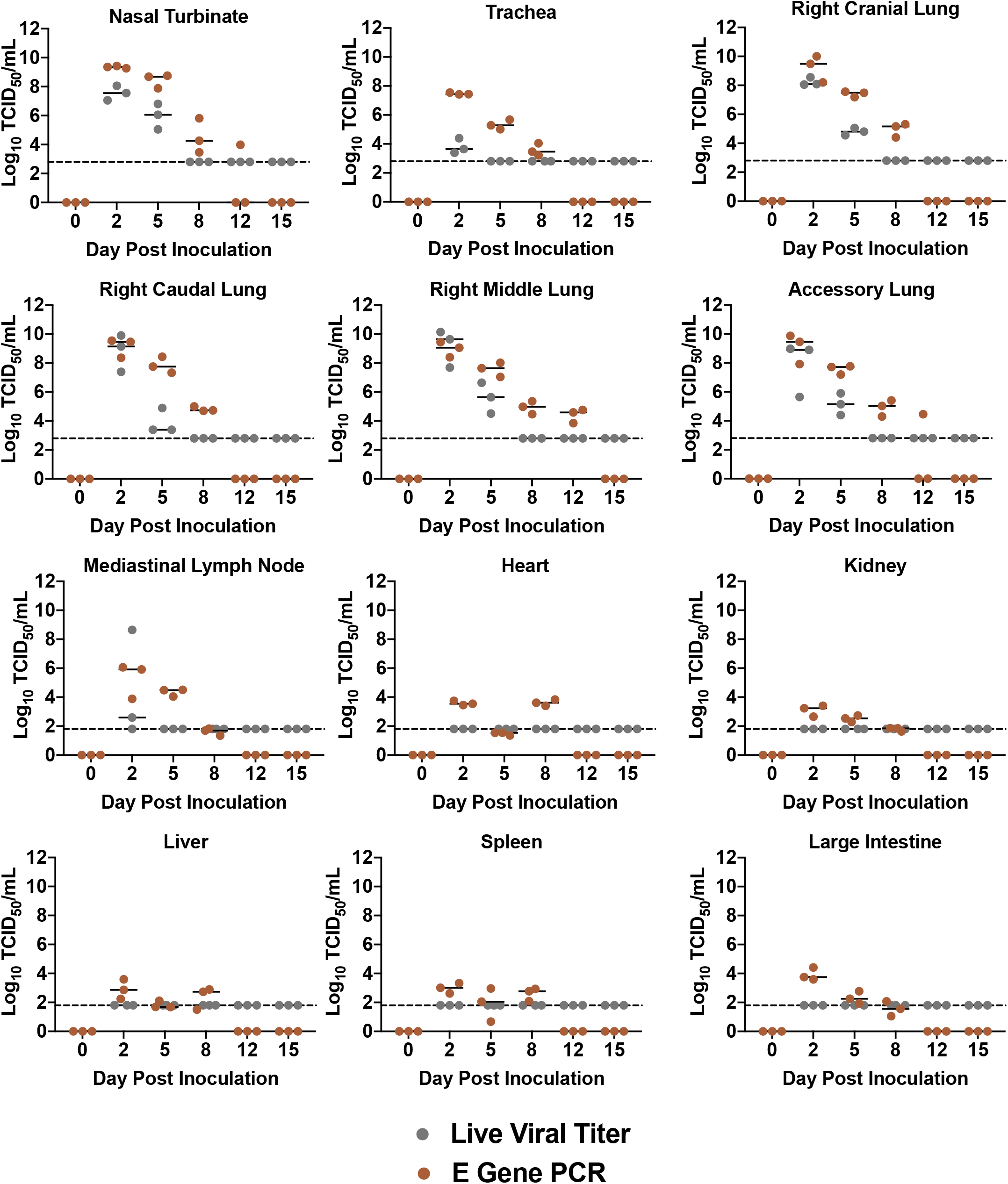
Live SARS-CoV-2 virus is confined to respiratory tissues and mediastinal lymph node while viral RNA is present in extrapulmonary organs. Animals were removed from the study on day 0 prior to infection as well as days 2, 5 8, 12 and 15 post inoculation. Respiratory tissues collected included the nasal turbinate, trachea, and lung (right cranial lobe, right middle lobe, right caudal lobe, accessory lung). Extrapulmonary organs analyzed included the mediastinal lymph node, heart, kidney, liver spleen and large intestine. All tissues were harvested to quantify virus by qPCR detection of the SARS-CoV-2 E gene (brown) and live viral load assay (grey). Infectious titer of tissues was determined by TCID50 virus titration assays and the TCID50/mL was calculated using the Reed and Muench method. Dotted line represents the limit of detection in the live viral load assay. Data points represent individual animals and line represents the average.

To determine if virus resolution was due to elicitation of neutralizing antibodies, we next quantified the neutralizing antibody titers in infected hamsters throughout the time course (**Figure 3A**). Plasma was isolated from blood collected at necropsy to be used for neutralization assays at 100 TCID50 per reaction of SARS-CoV-2. Neutralizing antibodies were detected as early as day 8 pi, at an endpoint titer of approximately 63 (geometric mean). This increased to a geometric mean of 160 on day 12 and peaked on day 15 at a geometric mean of 587. These observations suggested that hamsters are able to elicit high titers of neutralizing antibodies after SARS-CoV-2 infection. Nasal washes were collected throughout the study to determine viral shedding from the upper respiratory tract. vRNA was detected in the nasal washes up to 12 days after inoculation with live virus observed on days 2 and 5, peaking at day 2 at just over 6 TCID50/mL (Log10). We compared the live viral titer found in the nasal washes of all animals on day 5 to their respective weight loss on that day (**Figure 3B**). We found a Pearson correlation of −0.68 such that as the viral titer in the nasal washes increased, animals had increased likelihood of more severe weight loss (p-value= 0.00242).

**Figure 3.**
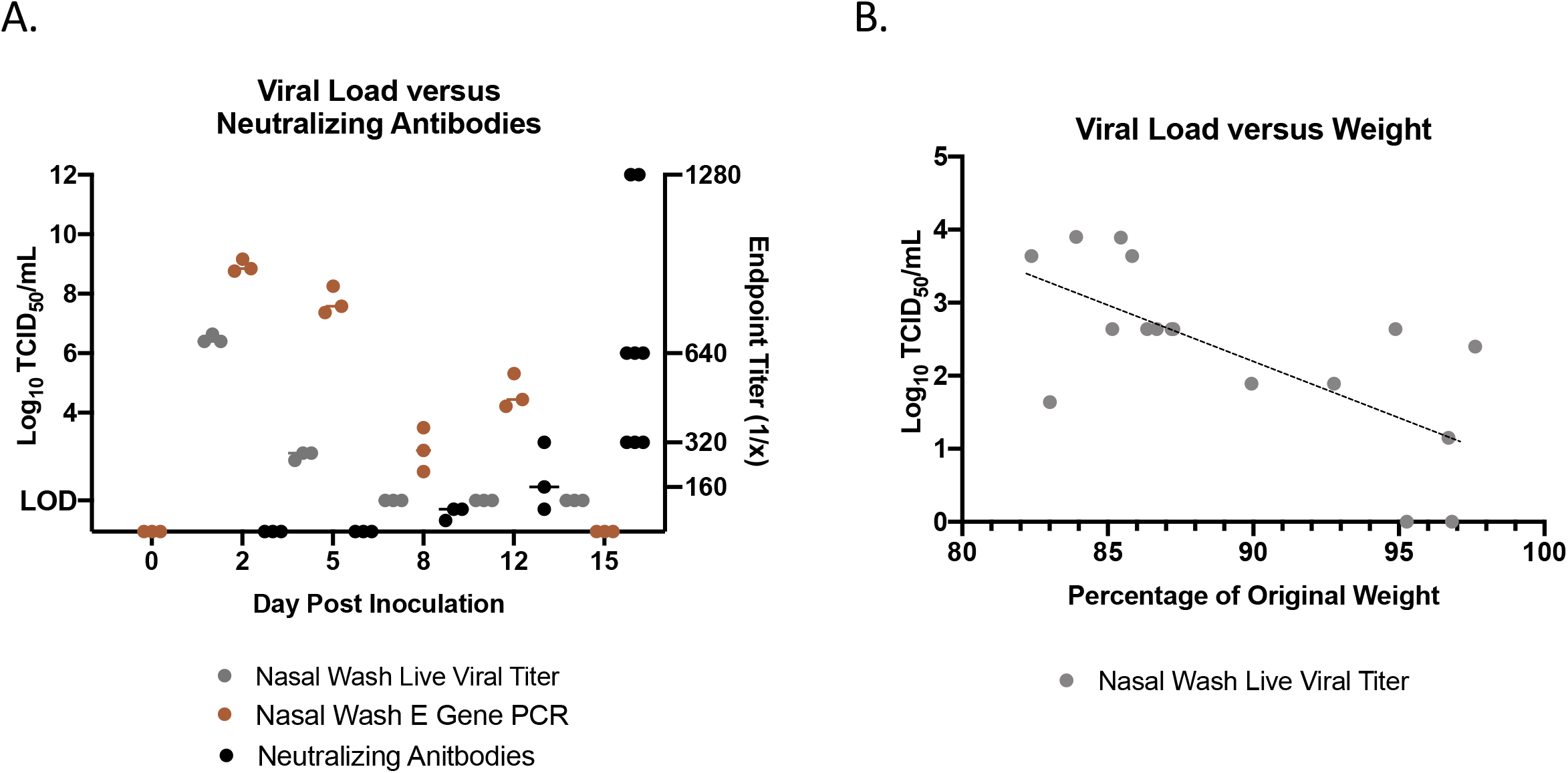
Viral shedding of SARS-CoV-2 increases with severity of clinical outcomes but diminishes as neutralizing antibodies are detectable in animal plasma. Nasal washes were collected to quantify virus by qPCR detection of the SARS-CoV-2 E gene (brown) and live viral load assay (grey). Infectious titers of the nasal washes were determined by tissue culture infectious dose titration assays and the resulting TCID50/mL was calculated using the Reed and Muench method. Virus neutralization titers were determined by standard neutralization assays using plasma collected throughout the time course (black) (A). Live viral titer in the nasal washes negatively correlated with weight loss on day 5 post infection. Pearson correlation=-0.68 (p-value=0.00242) (B).

Pneumonia and lung pathology are often the defining feature of respiratory viral infection, resulting in significant morbidity and mortality. To determine the extent of damage to the lungs in SARS-CoV-2 infected hamsters, and the ability of hamsters to recapitulate the severe respiratory involvement in human COVID-19 patients, we examined disease pathogenesis within the left lung of hamsters throughout course of infection (**Figure 4**). Microscopic analysis of H&E staining indicated an increase in damage in the lung over the course of the infection, with the most severe pathology noted on day 8 pi which partially resolved by day 15 (**Figure 4A**). Uninfected hamsters had clear alveolar space without evidence of inflammatory cell infiltration or hemorrhage. By day 2 pi, mononuclear cell infiltration was present in the alveolar and peribronchial spaces (yellow arrow). Cell infiltration continued on day 5 with increased infiltration. On day 8, the peribronchial space was characterized by the presence of inflammatory mononuclear cells. Epithelial cell sloughing in the bronchioles was noted (black arrow) and hemorrhage could be observed throughout the tissue (red arrow). Hemorrhage was resolved by day 12 although hyperplasia was evident (green arrow). By day 15 hyaline membrane formation (blue arrow) was observed with concomitant reduction of inflammatory cell infiltration.

**Figure 4.**
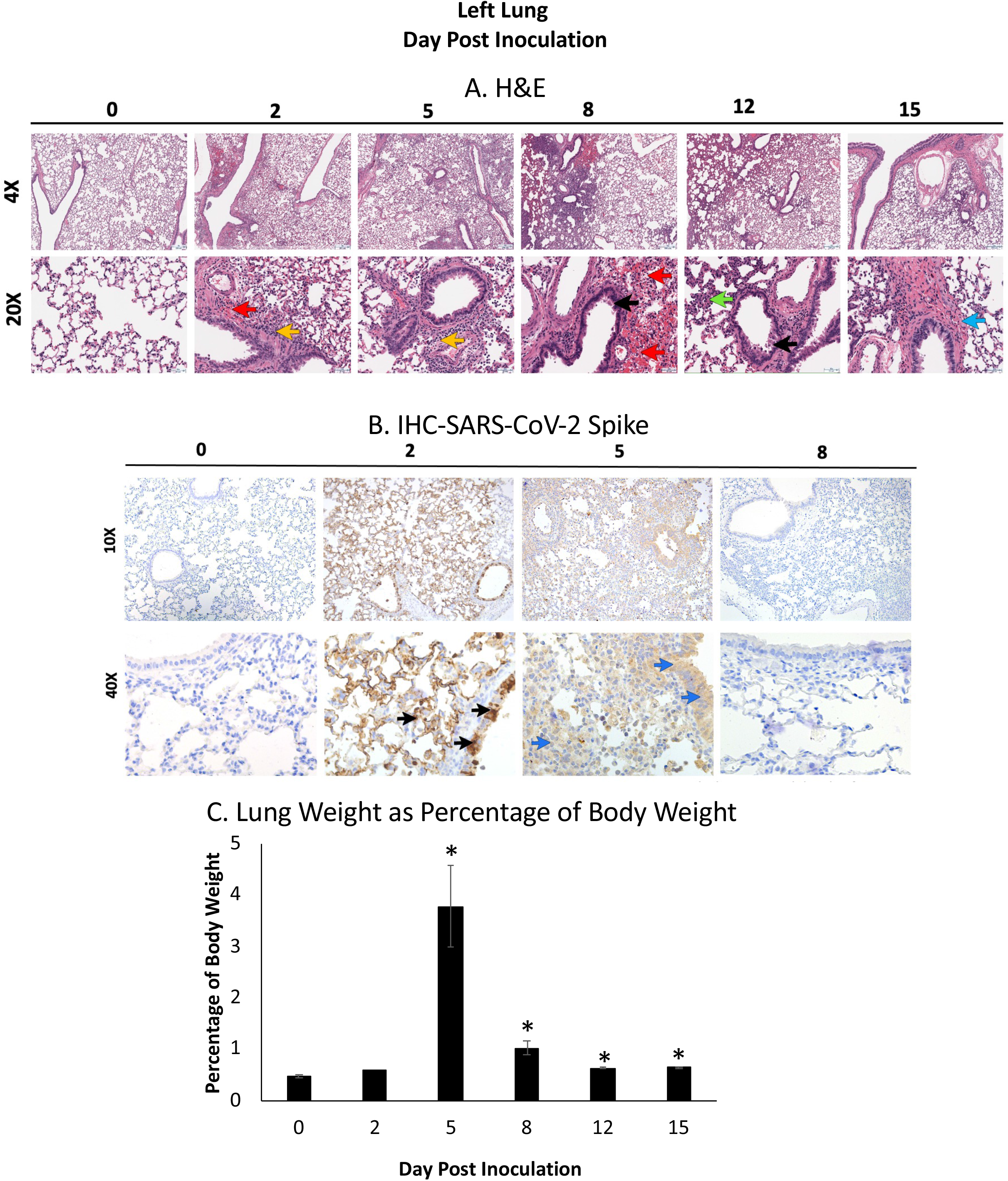
Inoculated hamsters have significant inflammation in the lung coinciding with SARS-CoV-2 Spike antigen staining. The left lung was collected from each animal at necropsy and perfused with 10% formalin prior to paraffin block embedding and H&E stained. Red arrows indicate hemorrhage; green arrows indicate hyperplasia; yellow arrows indicate mononuclear cell infiltration; black arrows represent cell sloughing; blue arrow represents hyaline membrane formation (A). The lung was stained for the presence of SARS-CoV-2 spike protein by IHC using an anti-spike rabbit monoclonal antibody (B). Black arrows indicate high amounts of viral antigen staining and blue arrows indicate the presence of low amounts of viral antigen staining. The entire lung was weighed at necropsy and lung gross pathology was represented as a percentage of total animal body weight (C). At least 3 animals per timepoint were analyzed. * indicates a p-value less <0.05 determined by ANOVA comparing hamsters on the days post inoculation to baseline (day 0). Error bars indicate +/- SE. Slides were visualized and imaged using the Leica DMI100 Brightfield microscope and DMC5400 20 MP color CMOS camera. Images were captured and 10X and 40X. Images shown are representative of 3 animals per group, per timepoint.

To validate viral load results in the lung, as well as visualize the location of the virus, we performed IHC staining of the left lung lobe to detect SARS-CoV-2 spike protein. No viral protein was detected in uninfected hamsters suggesting the specificity of the antibody used. On day 2 pi, the left lung had significant staining (black arrow) suggesting accumulation of SARS-CoV-2 spike antigen (**Figure 4B**). Staining was most prominent in the epithelial cells of the bronchiole, with diffuse staining throughout the epithelial cells of the alveolar wall. On day 5 light staining (blue arrows) was present through the airways and alveolar cells. No staining was detected on day 8 pi where the images appeared similar to images of uninfected bronchiole.

To quantify inflammation and inflammatory cell infiltration as well as edema in the lung, the weights of the lungs at necropsy were analyzed. The entire lung was weighed at necroscopy and a ratio to the animal’s body weight was calculated and then expressed as a percentage (**Figure 4C**). On day 0 and 2 pi, the lung was less than one percent of hamster body weight. This increased to a significant peak on day 5 at just under 4 percent of body weight on average. Lung weight decreased to above 1 percent and remained statistically significant from baseline throughout the remaining time course.

### Host gene expression and immunopathology profiling indicates a pronounced inflammatory response with blunted type I interferon signaling in the respiratory tract

Since the human data investigated systemic cytokine levels in the blood, we were interested in determining the immune responses at the site of infection as well as other affected organs. We investigated the levels of type I interferon response-related (IFN (Interferon)-beta, STAT (signal transducer and activator of transcription) 2, IRF (interferon regulatory factor) 1), type II interferon response-related (IFN-gamma, IRF2, STAT1, CXCL10), innate/adaptive mediators (IL-2, TGF (transforming growth factor)-beta, IL-21, IL-10, IL-4, IL-5, IL-13, IL-12, IL-1beta, IL-6, TNF), and other signaling molecules (PKR (protein kinase R), CCL20 and CCL22).

Human clinical reports have indicated that SARS-CoV-2 infection leads to a dysregulation of the interferon response^16^.We investigated the type I and II IFN responses by evaluating the expression of IFN-beta, STAT2, and IRF1 for the type I response and IFN-gamma, IRF2, STAT1, and CXCL10 for the type II response (**Figure 5**). IFN-beta was significantly downregulated in both the nasal turbinates and right cranial lung with the greatest inhibition on day 2. Downstream regulators of the type I IFN response were also decreased, including STAT2 and IRF1 which reached maximum inhibition later in the time course. We also examined the type II IFN response, as well as genes whose corresponding proteins act in both IFN pathways. IFN-gamma, the type II IFN, showed upregulation in the nasal turbinates, and most prominently in the lung (**Figure 5**). In both tissues there was an increase to a peak followed by resolution by day 15 pi. Nasal turbinate levels of IFN-gamma peaked on day 8 at a fold-change of 12, while levels in the lung peaked on day 5 at a fold-change of 17. In the respiratory tissues, type I IFN antagonist IRF2 showed significant increases, peaking on day 8 in the nasal turbinate and day 2 in the right cranial lung, at a fold-changes of 24 and 25, respectively. General IFN response molecules showed significant increases as well. STAT1 showed strong upregulation in both tissues on days 2 and 5, while CXCL10 peaked at day 2 in the nasal turbinates at a fold-change of close to 70. In the lung, CXCL10 peaked on day 8 at a fold-change of 30 but did not resolve back down to baseline and remained significantly increased relative to naïve animals on days 12 and 15 pi. From this data, it appears that while the type II IFN response remains intact, hamsters inoculated with SARS-CoV-2 show a dysregulated type I IFN response in respiratory tissues.

**Figure 5.**
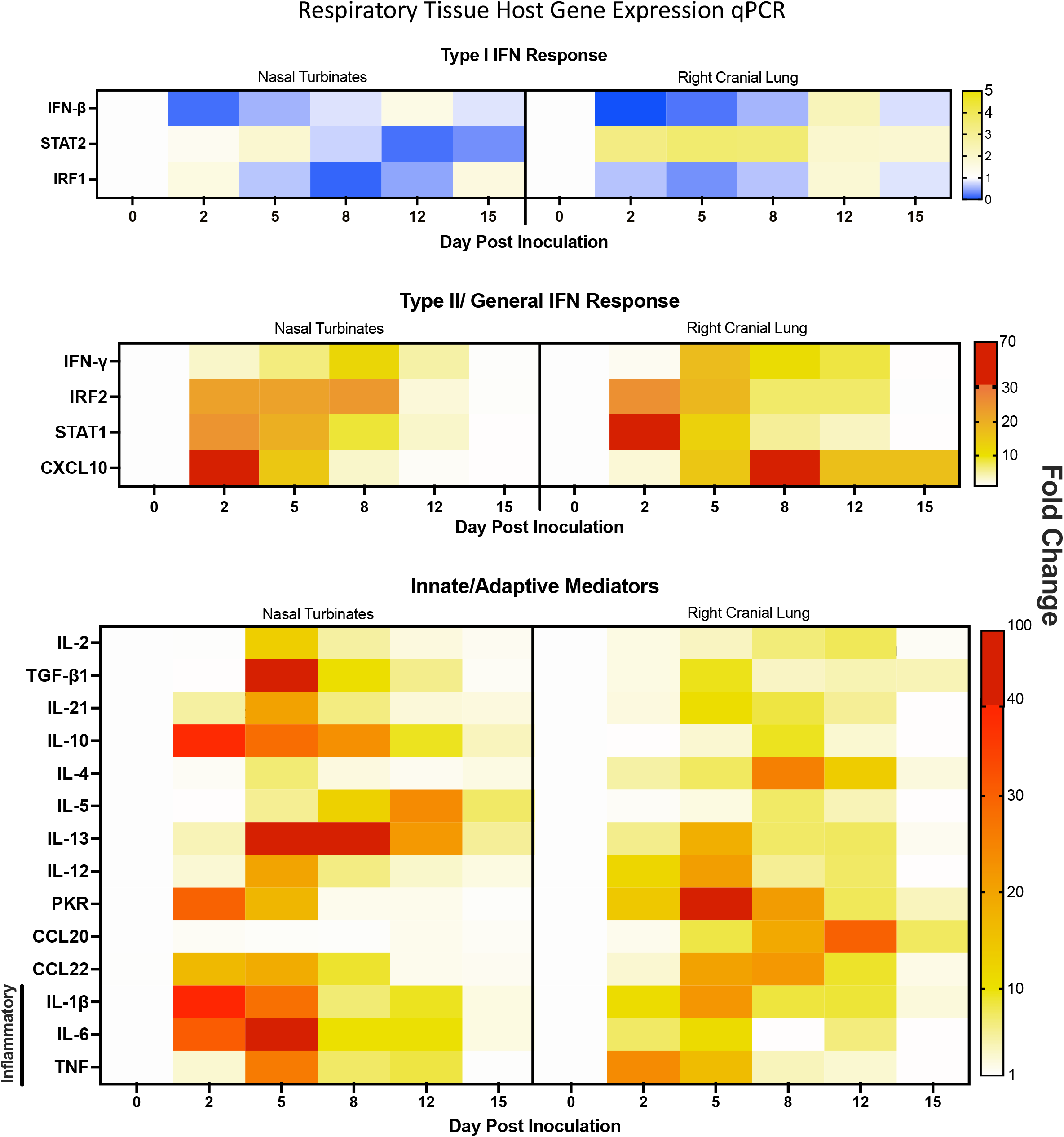
Gene expression analysis in the respiratory tissues of SARS-Cov-2 infected hamsters reveals significant interferon dysregulation with concomitant increases in inflammatory and immune mediators. qRT-PCR was performed on RNA extracted from nasal turbinate and right cranial lung tissue of SARS-CoV-2 inoculated hamsters. Samples were assessed for type I interferon response-related (IFN-beta, STAT2, IRF1), type II interferon response-related (IFN-gamma, IRF2, STAT1, CXCL10), and innate/adaptive mediators (IL-2, TGF-beta, IL-21, IL-10, IL-4, IL-5, IL-13, IL-12, IL-1beta, IL-6, TNF) using primers specific to hamster genes (**Table 1**). Fold-change was calculated via *ΔΔ*Ct against baseline (Day 0) with BACT as the housekeeping gene. At least three animals were analyzed for each timepoint.

**Table 1.**
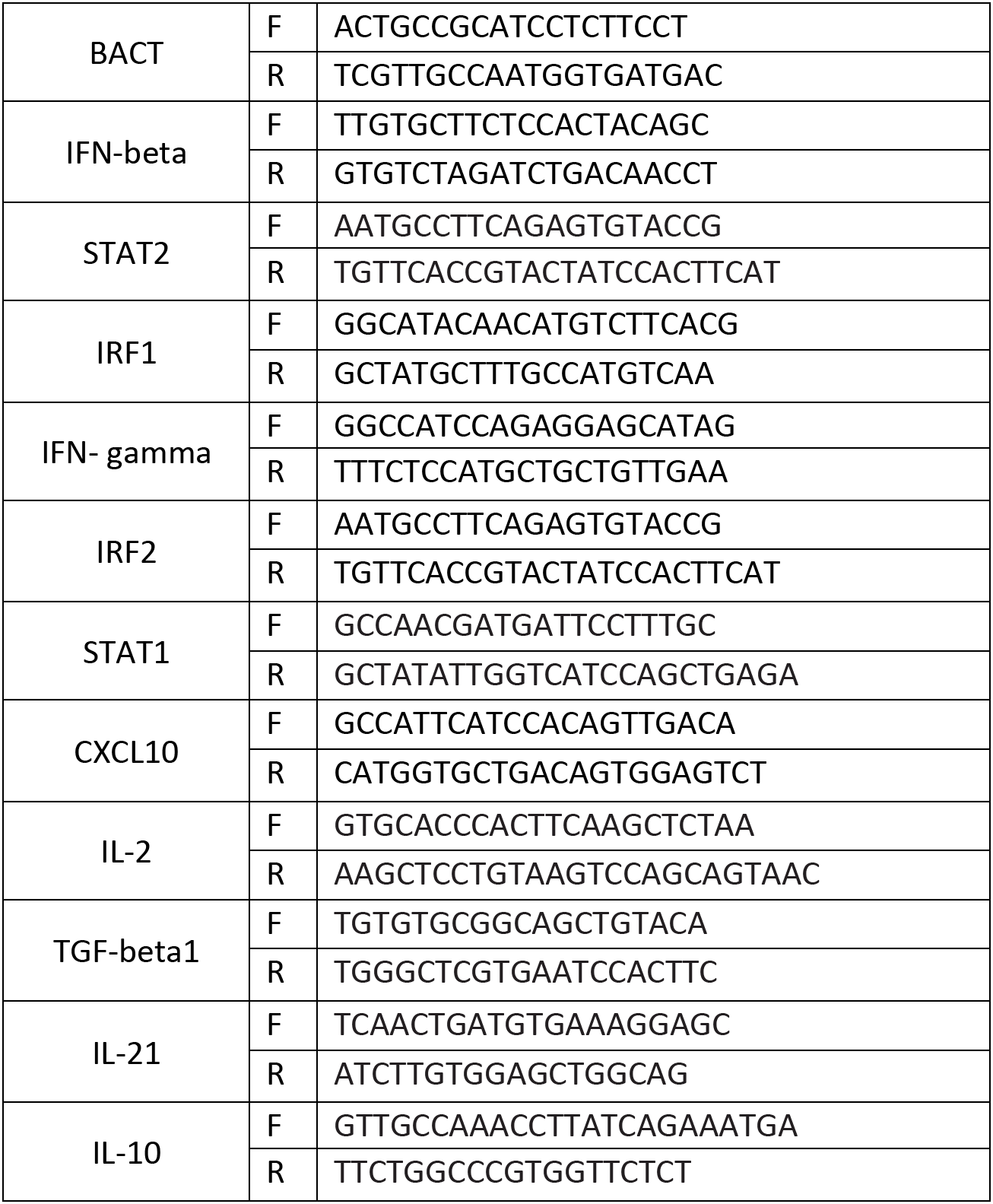
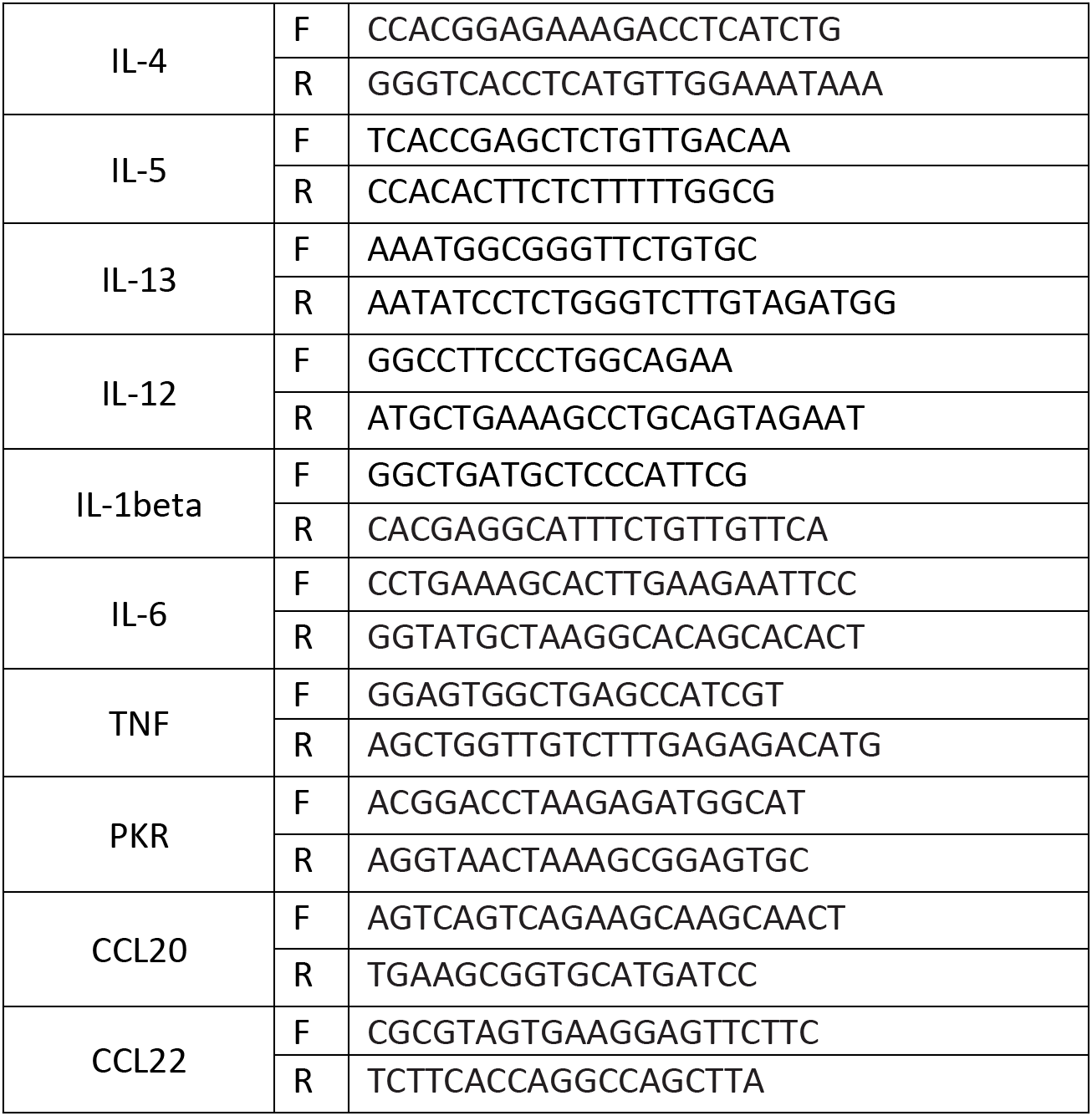
Syrian Hamster qRT-PCR Primers.

Additionally, we also assessed innate and adaptive gene regulation. Overall, these genes were upregulated in both the nasal turbinates and right cranial lung, with nasal turbinates typically showing higher (**Figure 5**). The nasal turbinates had peak gene expression of IL-2, TGF-beta1, IL-21, Il-4 and IL-13 on day 5 pi, with the highest expression coming from TGF-beta1 at a fold-change of 97, and IL-13 at 45. IL-10 expression peaked on day 2 at a fold-change of 38 but remained upregulated throughout the time course, significantly on days 5 and 10. IL-5 expression peaked on day 12 at a fold-change of 21. Within the lung, the highest expression per gene varied: TGF-beta1, IL-21 and IL-13 showed the highest upregulation on day 5; IL-10, IL-4 and IL-5 on day 8; and IL-2 on day 12. IL-4 and IL-13 showed the greatest upregulation, at 25 and 18 on their respective peak days. This data may suggest that an involved T cell response with the upregulation of Th2 cytokines (IL-4, IL-5 and IL-13), Th1 cytokines (shown previously IL-12 and IFN-gamma), as well as T regulatory and Th17 cytokines (IL-10 and TGF-beta1) point to an overall T cell response to SARS-CoV-2 infection. Therefore, we performed IHC to determine the presence of CD3+ and CD20+ cells within the respiratory tract (**Figure S2**). CD3+ staining was concentrated to blood vessels (red arrows) in the uninfected lung tissue with no staining within the alveolar spaces. The presence of CD3+ cells (**Figure S2A**, black arrows) increased in the alveolar space throughout the time course reaching maximum staining on day 8 post inoculation and then decreasing in intensity out to day 15. CD20 staining was not detected in uninfected tissue. Small numbers of CD20+ cells were detected on day 2, 5, and 8 post inoculation (**Figure S2B**, denoted by black arrows) which diminished on day 12 and 15 as indicated by grey arrows.

Analysis of the inflammatory markers within the nasal turbinates indicated significant transient increases of all inflammatory targets resolving by days 12 and 15. IL-1beta reached maximum levels on day 2 pi at a fold-change of 40 relative to day 0 uninfected animals (**Figure 5**). IL-6, IL-12 and TNF reached apex day 5 pi (fold-changes of 66, 20 and 26, respectively). Corresponding significance can be found in **Table S1**. In contrast, these transcripts reached apex between days 2 and 5, but at a lower fold-change compared to the lung. In both the nasal turbinates and in the lung several inflammatory cytokines remained elevated compared to baseline until day 12 pi. Taken together, the sites of infection showed a clear increase in inflammatory markers, especially IL-6 in the nasal turbinates and TNF in the lung.

### SARS-CoV-2 infection results in significant inflammation or immune regulation in the heart, lymph node, and spleen as well as eosinophil signatures in the large intestine

Immune markers of inflammation, adaptive immunity, and the antiviral responses were also assessed in non-respiratory tissues alongside histopathology of the tissues. We investigated the mediastinal lymph node, heart, kidney, spleen, liver and large intestine for regulation of immune targets categorized as above by type I IFN, type II/General IFN Response, and Innate/Adaptive Mediators (**Figure 6**) as well as for immunopathology (**Figure S1** and **Figure 7**). If initial assessment of histopathology suggested the involvement of specific immune mechanisms (**Figure S1**), high resolution scanning was performed to acquire a greater depth of information (**Figure 7**). Mediastinal lymph nodes were not able to be recovered on day 15. Corresponding significances for the gene expression can be found in **Table S2**.

**Figure 6.**
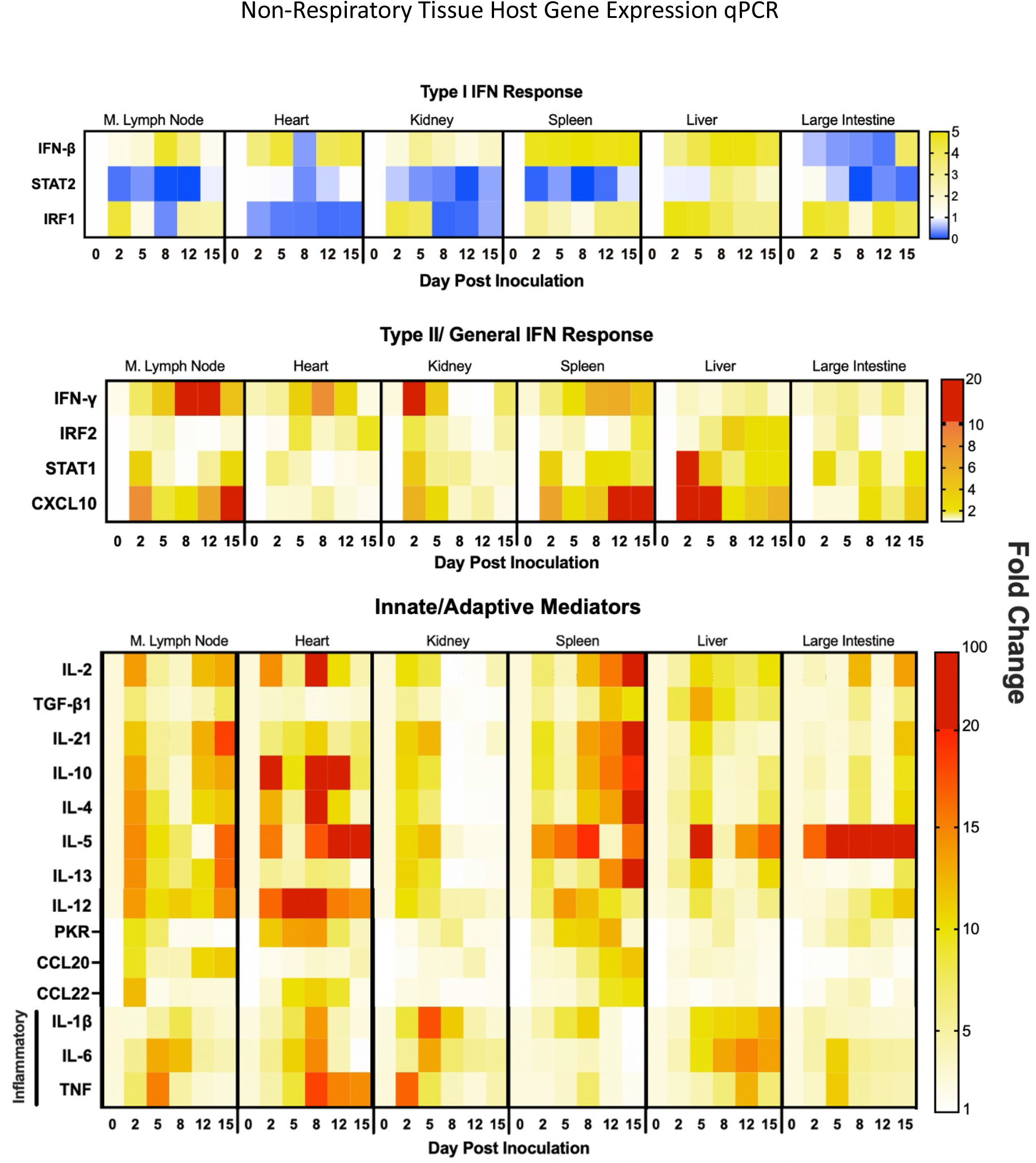
SARS-Cov-2 infection in hamsters leads to increases in inflammatory and immune mediator related genes in select extrapulmonary organs possibly indicating myocarditis, adaptive immune evolution, and eosinophil infiltration into the large intestines. qRT-PCR was performed on RNA extracted from mediastinal (M) lymph node, heart, kidney, spleen, liver and large intestine from SARS-CoV-2 inoculated hamsters. Samples were assessed for type I interferon response-related (IFN-beta, STAT2, IRF1), type II interferon response-related (IFN-gamma, IRF2, STAT1, CXCL10), and innate/adaptive mediator (IL-2, TGF-beta, IL-21, IL-10, IL-4, IL-5, IL-13, IL-12, IL-1beta, IL-6, TNF) genes leveraging PCR primers specific for hamster sequences (**Table 1**). Fold-change was calculated via *ΔΔ*Ct against baseline (Day 0) with BACT as the housekeeping gene. N is 3 for all timepoints.

**Figure 7.**
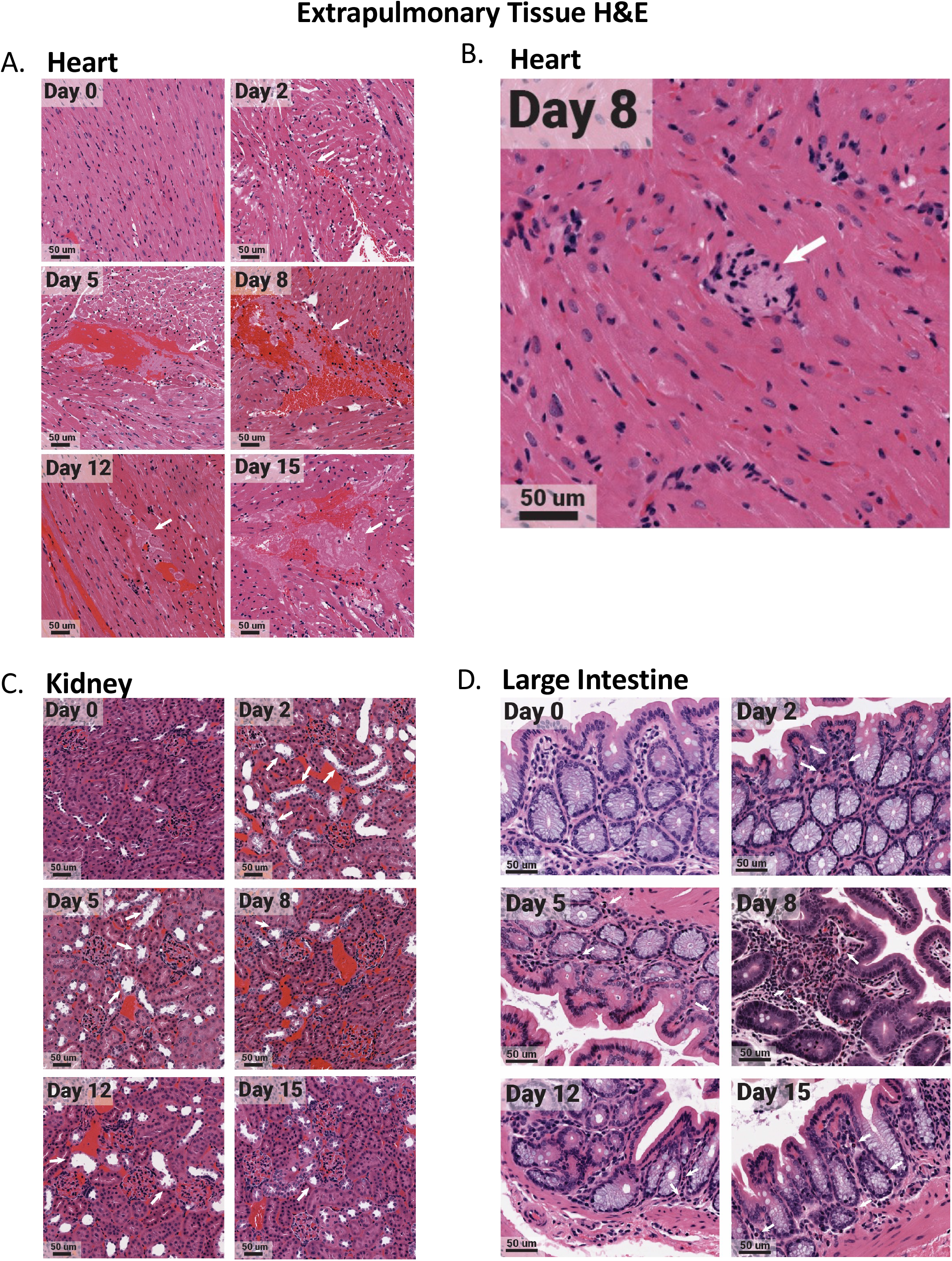
Select extrapulmonary organs have distinct pathology after SARS-CoV-2 infection with evidence of microthrombi and myocarditis in the heart, tubular damaging in the kidney, and eosinophil infiltration in the large intestine. The heart, kidney, and large intestine were collected from each animal at necropsy and fixed in 10% formalin prior to paraffin block embedding. Tissues were H&E stained. Heart tissues exhibited microthrombi (white arrows) over multiple days (A), as well as myocarditis on day 8 (white arrow) (B). The kidney had acute injury/tubular injury (white arrows) (C). The large intestine had eosinophil infiltration (white arrows) (D). Stained tissues were visualized and imaged using the Aperio ScanScope XT slide scanner. Images shown are representative of 3 animals per group, per timepoint.

The type I and type II IFN responses in all the extrapulmonary organs except the liver had similar regulation profiles as those of the lungs and nasal turbinates (**Figure 6**). Gene regulation was characterized by decreased type I responses and increases in type II responses. STAT2 was significantly decreased in the mediastinal lymph node, kidney, spleen, and large intestine. The heart had significant and sustained decreases in IRF-1 while the large intestine had decreases in IFN-beta (**Figure 6**). Conversely, marked increases in IFN-gamma was observed in the mediastinal lymph node and kidney and to a lower extent in the heart and spleen. CXCL10, the T cell chemoattractant, was increased to high levels in the spleen, liver, and mediastinal lymph node where the regulation was biphasic in the lymph node (**Figure 6**).

Regulation of the innate/adaptive mediator genes in the spleen and lymph node was seen for all immune targets examined with the exception of TGF-beta which was not effectively regulated. Specifically, significant regulation of IL-2, IL-21, IL-10, IL-4, IL-5, IL-13, and IL-12 was noted in these secondary immune organs. The regulation of the innate/adaptive immune mediators was particularly interesting in the heart where IL-2, IL-10, IL-4, and IL-5 were increased 100 times over baseline. The highest upregulation of inflammatory cytokines was detected in the heart followed by the kidney, liver, and mediastinal lymph node. In the heart, IL-12 was significantly increased early and, along with TNF, remained elevated at day 15 pi. Il-6 and IL-1beta were also transiently induced in the heart tissue on days 5 and 8 (**Figure 6**). Significant changes in the heart were observed by histological analysis (**Figure S1**). High-resolution imaging found evidence of microthrombi (**Figure 7A**, white arrows) as well as myocarditis (**Figure 7B**, white arrows) in the heart. IL-1beta and TNF were upregulated early in the kidney but decreased by day 8 to 12 pi (**Figure 6**) which coincided with infiltration of inflammatory cells into the kidney and possible acute kidney injury or tubular injury (indicated by the white arrows, **Figure 7C**). IL-6 expression was sustained in the liver where IL-1beta was also increased at late time points. The mediastinal lymph node had a biphasic increase in IL-12 with transient increases in IL-6 and TNF to fold levels of 12, 8 and 11, respectively. The large intestine had significant regulation of IL-5 with infiltration of eosinophils (**Figure 7D**), but minimal regulation of any other innate/adaptive mediators.

To accompany analysis of the extra-pulmonary organs, we analyzed blood chemistries in the animals over the time course to assess organ dysfunction (**Figure S3**). We analyzed glucose, albumin, alkaline phosphatase, alanine transferase, blood urea nitrogen, total bilirubin and total protein as a signs of liver function^33^. Calcium, potassium, albumin, amylase, blood urea nitrogen, and total protein were analyzed to assess the kidney^33^. Gastrointestinal health was evaluated through sodium, potassium, and alkaline phosphatase^33^. Cardiac function was measured using sodium, potassium and alanine transferase ^33^. Globulin upregulation was assessed as a sign of antigenic stimulation^33^. Of the 14 molecules investigated, only amylase and glucose were significantly increased in infected animals when compared to controls, peaking at 1700 U/L and 280 mg/dL, respectively, on day 8 pi. There was minimal regulation of albumin and total protein, markers of kidney and liver function. Other liver markers, including glucose (280 mg/dL), peaked later in the time course, including blood urea nitrogen on day 8 (22mg/dL) and alkaline phosphatase on day 12 (3 U/L). Alanine transferase, although peaking on day 2, was high on day 12 at just under 100 U/L. Kidney markers also peaked later, on day 8 for amylase (1700 U/L), and day 15 for potassium and blood urea nitrogen (12 and 22 mg/dl, respectively). Calcium, in contrast, peaked on day 2 at 2.6 mg/dl. Markers of gastrointestinal and cardiac health such as sodium, showed minimal regulation, while potassium increased relatively linearly throughout the infectious time course, peaking at day 15 (12 mg/dl). Alkaline phosphatase (gastrointestinal marker) was high on days 5 and 8 at 3u/L and alanine transferase (cardiac marker) was increased in infected hamsters relative to non-infected hamsters on days 2, 8 and 12 at 75-100 U/L. Lastly, globulin peaked on day 2 at 0.6 g/dL and decreased relatively linearly over the time course. Taken together, blood analysis indicates some dysregulation in extrapulmonary organs, as well as antigenic stimulation in response to SARS-CoV-2.

## Discussion

Originally SARS-CoV-2 infection was thought to mainly cause respiratory disease, but it is now clear that COVID-19 is a disease of many faces affecting several organ systems^31,32^. To gain a better understanding of the origin of disease and the myriad of clinical symptoms experienced, we investigated the immunopathology across the respiratory tract as well as non-respiratory organs during SARS-CoV-2 infection in Syrian hamsters. Infection with 10^5^ TCID_50_ of SARS-CoV-2 resulted in mild to moderate clinical illness. Infectious virus was mainly contained in the respiratory tract although viral RNA was present in all tissues evaluated. Host response profiling in the respiratory tract identified an inhibition of the type I IFN response with a concomitant increase in type II IFNs, inflammatory cytokines, innate and adaptive cytokines. Investigation of the extra-pulmonary organs also found a similar profile for type I and II IFN signaling; however, tissue-specific immune regulation was also identified. Specifically, significant expression of innate and adaptive mediators was identified in the heart and spleen. Since antiviral treatment during SARS-CoV-2 infection is only effective during a short window,^34^ severe COVID-19 is thought to be a consequence of a dysregulated immune response^35^. Understanding immune pathogenesis is essential for identifying therapeutic targets and treating acute non-respiratory symptoms associated with COVID-19 including heart palpitations or migraines and longer-lasting conditions in the recovery phase such as Post-Acute Sequelae of SARS-CoV-2 infection (PASC), also referred to as long-COVID^36^.

The primary site of infection influencing transmission for SARS-CoV-2 in people is the respiratory tract. Therefore, the initial focus of our investigation was the respiratory tract, including associated immune responses and immunopathology. Here we have demonstrated both upper and lower respiratory infection and associated disease following SARS-CoV-2 infection in Syrian hamsters. Viral infection in the lung led to severe pathology characterized by infiltration of inflammatory cells resulting in bronchopneumonia by day 5 pi and hemorrhage by day 8. These findings were consistent with SARS-CoV-2 infection in prior Syrian hamster studies^21,22,37,38^ and larger animal models including rhesus macaques^39^ as well as the severe pneumonia identified in some human COVID-19 patients^40–42^. The consistency of the data supports the use of the hamster to model SARS-CoV-2 infection human respiratory disease and the associated immunopathology. To this end, we identified the upregulation of key inflammatory genes, such as IL-6 and TNF in the lungs and nasal turbinates confirming previous immune modeling in hamsters^38,43^. These findings are important as these inflammatory markers have been shown to be predictors of severe disease via assessment of blood in humans^44–48^. More in-depth analysis of the host response to SARS-CoV-2 in the respiratory tissues of hamsters has been done using transcriptome sequencing analysis^49,50^. Hoagland and colleagues researchers assessed the lungs of SARS-CoV-2 infected hamsters for host immune gene regulation and found a prominent upregulation of IL-6, TNF, and IL-1beta^49^ consistent with our PCR results. In another study single cell sequencing analysis revealed a prominent T cell response including upregulation of IFN-gamma and IL-2^50^, two markers we also noted to be highly expressed in the lungs. Importantly, the qRT-PCR analysis we reported in our present study is in agreement with these other studies thereby validating the use of qRT-PCR as a method to quickly assess the effectiveness of vaccines and therapeutics being evaluated in the hamster SARS-CoV-2 model. Our qRT-PCR profiles provide targeted biomarkers that can easily be employed.

The antiviral response is essential for the initial control of a virus by active destruction of viral replication and immune stimulation to limit the spread to other cells. After identifying the presence of a virus or viral antigens, the induction of interferons is central to the establishment of an antimicrobial state within cells where the balance between the interferons and other innate cytokines is critical to effective action^51^. Recently, inhibition of type I interferons (IFN-α and IFN-beta) has been associated with severe disease in COVID-19 suggesting the virus is able to block antiviral immunity leading to increased pathogenesis ^16,52,53^. In our study, we found that hamsters infected with SARS-CoV-2 have a prototypical type II IFN response. In contrast, the type I IFN response is downregulated, as evidenced by the decrease in IFN-beta gene expression and increase in the IRF2 gene expression, a type I antagonist. Numerous studies have demonstrated impaired type I IFN responses following SARS-CoV-2 infection *in vitro* ^16,54,55–57^. The IFN dysregulation we report here has not yet been analysed in the respiratory tissue of humans with severe COVID-19, but systemic immune analyses of blood have indicated similar results ^16,52,53,58^. Although the mechanisms inhibiting type I interferon responses have not been fully elucidated, viral and host factors have been implicated in this dysregulation ^59,60^.

Dysregulation of the interferon response is a strategy employed by viruses to evade host immunity. Previous investigations of the host response to SARS-CoV-1 (severe acute respiratory syndrome coronavirus 1) and MERS-CoV (Middle East respiratory syndrome coronavirus) infection suggest that the severe coronaviruses employ this strategy^61–63,64,65^. The 2020 study by Xia et al. determined this dysregulation utilized by the highly pathogenic coronaviruses extends to SARS-CoV-2 by demonstrating that all three of these beta-coronaviruses downregulate the type I interferon pathway. Specifically, NSP1, NSP6 and NSP13 ^56^ interfere with IFN-beta translation, either by blocking host ribosome in the case of NSP1^66^ or targeting upstream proteins necessary for IFN-beta induction in the case of NSP13^67^. Other SARS-CoV-2 virus proteins have also been implicated in interferon downregulation^68^. ORF3b and ORF9b both specifically target a mitochondrial antiviral-signaling protein complex in order to inhibit type I IFN siglalling^69,70^. Host factors affecting SARS-CoV-2 infection outcomes include age and sex; furthermore, comorbidities have also been implicated in interferon dysregulation^71^. A 2020 study by Trouillet-Assant et al. examined COVID-19 patient outcomes in a cohort including individuals systemically expressing type I IFN and individuals who had significantly lower levels of both IFN-alpha and beta. They found patients lacking type I interferon expression were older, and more likely to be male compared to those who did express type I IFNs ^72^. In a similar study significantly higher systemic levels of IFN-alpha2 were identified in females compared to levels in males^73^. In our recent study, we found that older adult male ferrets had a delayed type I interferon response when compared to adult females, with males lacking induction of essential antiviral genes including OASL, MIX1 and IDG15 which coincided with increased viral burden^74^. In our present study using the hamster model, all of our animals were male suggesting that investigation into sex-biases are needed to gain a full understanding of our observed type I interferon dysregulation. At this time, it is not known why or how host factors influence interferon responses to SARS-CoV-2. A study by Bastard et al. identified the presence of autoantibodies against type I interferons in males with severe COVID-19 ^75^. This offers a possible explanation of the immune mechanisms involved in decreased type I interferon responses in a subset of human patients. The mechanism for reduced interferon responses in our present study, where all hamsters were adult male animals, awaits further investigation. Moreover, taken together, the immune profile we have identified together with the respiratory tract pathology suggests hamsters to be an ideal model to explore these mechanisms and test therapeutics such as type I interferons^54,55,76^ or interferon lambda^77,78^, both of which have demonstrated effectiveness in treating COVID-19 patients^79–81^ but more research is needed.

COVID-19 is a disease that involves the pathogenesis of both respiratory and non-respiratory organs ^31,32^, however little is understood about the host response outside of the respiratory tract. Although we found upregulation of inflammatory and type II interferon genes in all non-pulmonary tissue, the most striking response was the increase of immune modulatory cytokines in the heart. Within the heart our analysis identified the increase of cytokines including IL-10, IL-2, IL-4, IL-5, and IL-12, where some cytokine mRNA transcripts were increased 100-fold. Although these cytokines can be produced by different T cell subsets, the absence of TGF-beta suggests the presence of T-regulatory cells less likely. Additionally, the absence of IL-13 decreases the likelihood of a TH2 response. Apart from T-lymphocytes, the increase in gene expression of IL-10, TNF, and IL-6 could be attributed to monocytes or macrophages^82^. The expression of IL-5 and IL-4 also suggests the presence of eosinophils^83^. A large number of clinical studies have now detected cardiac involvement in COVID-19^84–90^. In fact, it is estimated that approximately 30% of people who have recovered from COVID-19 have some type of cardiac injury ^91^. A German study of 100 COVID-19 patients found that myocardial inflammation was common even months after recovery from SARS-CoV-2 infection^90^. Within the tested cohort, 78% of patients experienced cardiac-related symptoms regardless of their degree of COVID-19 severity^90^. Monocytes are among cell types known to infiltrate the heart in myocardial inflammation, producing IL-10, IL-1 beta, IL-6 and TNF^92^, as well as IL-12^93,94^, all of which were upregulated in the heart post SARS-CoV-2 inoculation in our present study. In addition, another subset of cardiac issues related to COVID-19 have been attributed eosinophils resulting in eosinophilic myocarditis^95^. Other viruses are known to cause eosinophilic myocarditis after infection^96–99^. This pathology typically results from the virus infecting the tissues of the heart for instance with Coxsackie virus B3 infection^100^. Eosinophilic myocarditis has been diagnosed in COVID-19 patients, where upon autopsy the heart tissue exhibited infilation of inflammatory cells including lymphocytes, macrophages and prominently, eosinophils^95^. In this case, the increase in eosinophils and overall inflammation in the heart was found despite the lack of live virus in the heart at the time of death^101^. We found evidence in our study of eosinophilic myocarditis in the histopathology and also the cytokine signature found in the heart. It is possible in our study that eosinophils lead to inflammatory mediated cardiomyopathy via upregulation of IL-4, which was increased in terms of gene expression in the hearts of hamsters on day 8 post SARS-CoV-2 inoculation. Increase of potassium and slight increases in alanine transferase in the plasma of infected hamsters also may have supported cardiac dysregulation. With this in mind, the mechanisms regulating cardiac injury related to COVID-19 are still not known. Considering the prevalence of cardiac involvement in COVID-19 patients, more research is needed to understand the mechanisms of disease. Here we have provided evidence that hamsters may be useful for understanding the defining mechanisms of cardiac injury during SARS-CoV-2 infection and evaluating therapeutics for this pathology.

Immunologically, our analyses in the spleen and mediastinal lymph nodes suggest the induction of adaptive immunity. The mediastinal lymph node identified a notable biphasic regulation in T cell related genes, with upregulation in the gene expression of IL-2, IL-21, IL-10, IL-4, IL-5 and IL-13 on days 2 and 15. This profile suggests early antigen presentation within the draining lymph node shortly after lung infection by dendritic cells, which would be supported by the day 2 upregulation of IL-12 production in the lymph node^104,105^. Dendritic cell migration leading to T cell activation eventually culminates in IL-2 stimulating T cell growth and proliferation ^106^. Globulin levels in the blood on day 2 post infection in infected hamsters also support early antigenic stimulation. The upregulation of IL-4, IL-5, and IL-13 suggests a TH2 skewed response ^107^. Early studies have demonstrated that asymptomatic or mild COVID-19 cases with immune skewing towards a systemic TH2 characterized by increases in IL-4 and IL-10^108^. IL-10, a regulatory cytokine, was also upregulated on day 2 in our study. Our data suggests that activated T cells are recruited to tissues with live viral infection such as the nasal turbinates, where we observed an increase in T cell related genes on day 5 post inoculation. The secondary upregulation of genes in the lymph node on day 15 also suggests an expansion of T cell subsets. In the absence of infectious virus at this time, this could be an increase in T cell subsets that directly contribute to the development of a memory response. As central memory T cells also produce IL-2^109^ our data showing increased IL-2 on day 15 post inoculation may be indicative of T cell memory. IL-21 was the most upregulated T cell related gene on day 15. IL-21 is primarily produced by T-follicular helper cells, a subset that aids germinal center B cells ^110^, particularly skewing B cells toward affinity maturation and memory development. A 2021 study by Lakshmanappa et al. assessed SARS-CoV-2 infection in rhesus macaques, revealing a robust T cell response in the mediastinal lymph node with specific increases in CD4, as well as T follicular helper cell markers CD28 and CXCR5^111^. It is possible the increased T cell gene expression observed in our study is also indicative of a germinal center response. An elevated systemic T cell response has also been observed in patients with SARS-CoV-2 who recover and develop long-term memory ^105,112^. A deeper analysis with increased markers of germinal center reactions for lymph node assessment would be beneficial for vaccine studies.

The elevated lymph node activity could also be indicative of pathogenesis. The lymph node was the only non-respiratory organ where live virus was detected. Live viral infection of the lymph node, as well as the spleen, has been observed in humans who succumbed to SARS-CoV-2 ^113^. In a study by Kaneko et al., human lymph node and spleen samples from deceased COVID-19 patients lacked BCL6+ B cells as well as entire germinal centers in general. The authors hypothesized this was the cause of poor humoral response to SARS-CoV-2^114^. Since we identified a robust neutralizing antibody response that was detected by day 8 post infection which coincided with a decrease in viral shedding, it is more likely that the live virus found in the lymph nodes of our study were indicative of functional B cell responses and germinal center reactions.

The spleen also showed prominent immune responses characterized by increases in gene expression linearly from day 8 to day 15. This late upregulation correlates with typical splenic involvement with the immune response to pathogens which would occur after activation in local draining nodes ^115^. The spleen, a lymphoid organ does also activates T and B cells in response to viral antigen. High gene expression of IL-2 and IL-21 is suggestive of T-follicular cell as well as germinal center dependant B cell activity, as we saw in the lymph node ^116^. Additionally, the response in the spleen may be a consequence of viremia, where plasmacytoid dendritic cells (pDCs) are involved in viral antigen recognition via TLR7 and TLR9 for CD8+ T cell and Natural Killer T (NKT) cell activation. Both of these cell types function in the clearing of virally infected cells after activation in the spleen by IFN-beta and IL-12^117^, which were increased in our model. IL-12 and IL-10 were upregulated in the spleen of infected hamsters, potentially pointing to macrophage phagocytic activity^118,119^. Also, innate lymphoid cells (ILCs) in the spleen support tissue repair in response to injury and produce IL-5 and IL-13^120^. Taken together, this later response in the spleen points to the recycling function of the spleen, as well as potential lymphocyte activation for long-term memory responses.

The large intestine showed minimal regulation of the assessed cytokines, with the notable exception of IL-5. IL-5 gene expression increased significantly on days 2 through 15 with a peak fold-change of 74 relative to baseline on day 8. IL-5 is predominantly secreted by type 2 T-helper cells and eosinophils^121^. Given the lower gene expression levels of other Th2 related genes, such as IL-4 and IL-13, it could be that IL-5 is being produced primarily by eosinophils which can been seen in our histological analysis of the large intestine. Eosinophils play a direct role in antiviral defence, as evidenced by RNAses in their granules and have been associated with the clearance of other respiratory viruses^122,123^ Large numbers of eosinophils are also found in the gastrointestinal tract^124^. In our study, the large intestine of infected hamsters did not exhibit live virus. The large intestine was, however, positive for SARS-CoV-2 viral RNA on days 2 and 5 post inoculation, a finding consistent with Sia et al., who also observed vRNA in the colon in the absence of an inflammatory response^22^. Up to 20% of COVID-19 patients report gastrointestinal symptoms such as diarrhea^45,125^ with virus being shed in the feces^126^. This could be explained by the decrease in eosinophils that has been reported in cases of severe COVID-19^127–129^. The increase in IL-5 gene expression by eosinophils could contribute to the antiviral state of the large intestine, thereby preventing infection. Further investigation is needed to confirm the exact cell type leading to this heightened IL-5 response.

Our study offers important insights into SARS-CoV-2 infection and COVID-19 disease regulation. Preclinical models are the cornerstone of biomedical research offering the ability to experimentally isolate disease mechanisms and also evaluate potential medical countermeasures including immunomodulatory drugs, antivirals, and vaccines. As SARS-CoV-2 first emerged, we had limited understanding of the mechanisms of pathogenesis, no identified therapeutics or prophylactics, and no preclinical model to leverage for medical countermeasure discovery. Since the time of viral emergence, huge leaps have been made especially in preclinical model development, but our understanding of COVID-19 and its many manifestations is still limited. Our study confirmed the viral tropism and respiratory pathology previously identified in our SARS-COV-2 hamster studies but was also able to extend our understanding of immune regulation and pathogenesis within the respiratory tract and beyond.

## Methods

### Ethics Statement

All work was conducted in accordance with the Canadian Council of Animal Care (CCAC) guidelines, AUP number 20200016 by the University Animal Care Committee (UACC) Animal Research Ethics Board at the University of Saskatchewan. Hamster procedures were performed under 5% isoflurane anesthesia.

### Virus

The SARS-CoV-2 isolate /Canada/ON/VIDO-01/2020 was used for infections and *in vitro* assays. This virus was isolated from a patient at a Toronto hospital who had returned from Wuhan, China^28^. The second passage viral stock was sequenced (GISAID – EPI_ISL_425177) to confirm stability of the SARS-CoV-2 virus after culture in vDMEM (DMEM (Dulbecco’s Modified Eagle Medium) (*Wisent Bioproducts (Cat # 319-005-CL)*), 2% fetal calf serum (*Wisent Bioproducts (Cat # 090-150)*), 5 mL 100x penicillin (10,000 U/mL)/streptomycin (10,000 μg/mL), and 2 μg/mL TPCK-trypsin) on Vero-76 cells. All work with infectious SARS-CoV-2 virus was performed in the Vaccine and Infectious Disease Organization’s (VIDO) Containment Level 3 (CL3) facility (InterVac) in Saskatoon, Saskatchewan, Canada.

### Animals, Infections, and Tissue Collection

Male hamsters aged 6-8 weeks were purchased from Charles River Laboratories (Wilmington, USA). Hamsters were anesthetized with isoflurane for intranasal inoculation with SARS-CoV-2 at 10^5^ TCID_50_ per animal. Animals were euthanized on days 2, 5, 8, 12, and 15 post inoculation, and tissues were collected for virological, immunological, and pathological analysis. Animal weight and temperature were monitored throughout the course of the investigation. Weight and temperature were calculated as a percentage of original values from day 0. Blood was collected in BD Vacutainer^®^ EDTA coated blood collection tubes for plasma separation.

### Viral Titers

Nasal washes (0.3 mL) were collected post inoculation in anesthetized hamsters. Tissues collected at necropsy were homogenized in serum free DMEM using a Qiagen TissueLyzer. A 1:10 dilution series of sample was established in viral growth media (vDMEM) for TCID_50_ virus infection titration assays. Viral load was based on cytopathic effect (CPE) observed on day 5 after cell infection and used to calculate the 50% endpoint using the Reed-Muench method.

### Viral Neutralization Assay

Plasma was heat-inactivated at 56°C for 30 m and then serially diluted 1:2. Virus was diluted to 100 TCID_50_ per well in vDMEM and used at a 1:1 ratio to plasma, incubated at 37°C for 1 h, and added to cultured Vero-76 cells in 96-well plates for 1 h at 37°C. Endpoint neutralization titer was based on inhibition of CPE observed on day 5 after cell infection.

### RNA extraction and quantitative Real-Time PCR (qRT-PCR)

Tissue RNA was extracted using the Qiagen© RNeasy Mini kit cat. # 74106 (Qiagen, Toronto, Canada) according to the manufacturer’s instructions. vRNA was extracted from nasal washes using the Qiagen© QIAamp Viral RNA Mini Kit cat. # 52904 (Qiagen). All host qRT-PCR was performed in triplicate on cDNA synthesized as previously described ^29^ (Table 1). vRNA was quantified by Qiagen© Quanti-fast RT probe master mix (Qiagen, Toronto, Canada) using primer/probe sets specific for the SARS-CoV-2 E gene. The reactions were performed on a StepOnePlus^TM^ Real-Time PCR System in a 96-well plate (Thermo Fisher) as previously described^30^.

### Clinical Chemistry

Blood collected at necropsy was centrifuged to yield plasma. Blood biochemistry analyses were performed on 100 µL of plasma using the VetScan VS2 instrument (Abaxis Veterinary Diagnostics). Sample was loaded into the VetScan ® Comprehensive Diagnostic Profile Rotor. The rotor was loaded in the VetScan VS2 Analyzer and analyzed as plasma samples from male hamsters. Results were analyzed in Prism 9.

### Histopathology and immunohistochemistry

The tissues collected for histopathology were submersed in formalin for at least 7 days within the CL3 lab. Formalin-fixed tissues were paraffin-embedded, sectioned, slide-mounted, and stained at Prairie Diagnostic Services (Saskatoon, Saskatchewan). Tissue samples were stained with hematoxylin and eosin (H&E) or immunostained to detect proteins of interest. H&E stained tissues were imaged first using a Leica DMI100 Brightfield Microscope and DMC5400 20 MP color CMOS camera (Leica Microsystems, Concord, Ontario, Canada). Further imaging was done using a Aperio ScanScope XT on select tissues, as indicated (Leica Biosystems, Nußloch, Germany). To detect the SARS-CoV-2 antigen, the VIDO polyclonal rabbit anti-SARS-CoV-2 AS20-014 was used (1:400 dilution). Immunohistochemical staining for host antigen was conducted using an automated slide stainer (Autostainer Plus, Agilent Technologies Canada Inc., Mississauga, ON). Epitope retrieval was performed in a Tris/EDTA pH 9 buffer at 97°C for 20 minutes. The CD3 (Rat anti-CD3 clone CD3-12, P. Moore, UC-Davis, Davis, CA) and CD20 (Rabbit anti-CD20, Thermo Fisher Scientific, Waltham, MA) primary antibodies were applied for 30 m at 1:25 and 1:400 dilutions, respectively. Binding of the primary antibodies was detected using an HRP-labelled polymer detection reagent. An HRP-labelled polymer detection reagent (EnVision+ System - HRP Labelled Polymer, Agilent Technologies Canada Inc., Mississauga, ON) was used to visualize viral and host protein presence. Immunostained tissues were all visualized, and images were captured using a Leica DMI100 Brightfield Microscope and DMC5400 20 MP color CMOS camera (Leica Microsystems, Concord, Ontario, Canada).

### Statistical Analysis

Unpaired, unequal variance, two-tail Student’s *t*-test or one-way ANOVAs were conducted using GraphPad Prism8 (San Diego, USA). A *p* value of ≤ 0.05 was considered statistically significant.

## Data Availability

The data that support the findings of this study are available from the corresponding author upon reasonable request.

## Acknowledgments

We greatly acknowledge the review of this manuscript by Dr. Kristen Kindrachuk and the assistance of Swarali Kulkarni and Dr. Vinoth Manoharan.

## Funding

A. Kelvin is funded by the Canadian 2019 Novel Coronavirus (COVID-19) Rapid Research Funding initiative the Canadian Institutes of Health Research (CIHR) (grant numbers OV5-170349, VRI-172779, and OV2 – 170357) and Atlantic Genome/Genome Canada, Scotiabank COVID-19 IMPACT grant, and the Nova Scotia COVID-19 Health Research Coalition. D. Falzarano is funded by the Canadian Institutes of Health Research (CIHR), grant number OV5-170349. J. Kindrachuk is funded by a Tier 2 Canada Research Chair in the Molecular Pathogenesis of Emerging and Re-Emerging Viruses provided by the Canadian Institutes of Health Research (Grant no. 950-231498). This article is published with the permission of the Director of VIDO. VIDO receives operational funding from the Canada Foundation for Innovation through the Major Science Initiatives Fund and by Government of Saskatchewan through Innovation Saskatchewan and the Ministry of Agriculture.

## Contributions

Conceptualization: AAK, JK, VG; Investigation: AAK, JK, MEF, AK, UG, YL, RR, CS, SM; Analysis: AAK, DF, MEF, AK, CS, UG, SM; Writing: AAK, MEF, MR, SM, VG

## Competing Interests

The authors declare no competing interests.

## Supplementary Tables

**Table S1.**
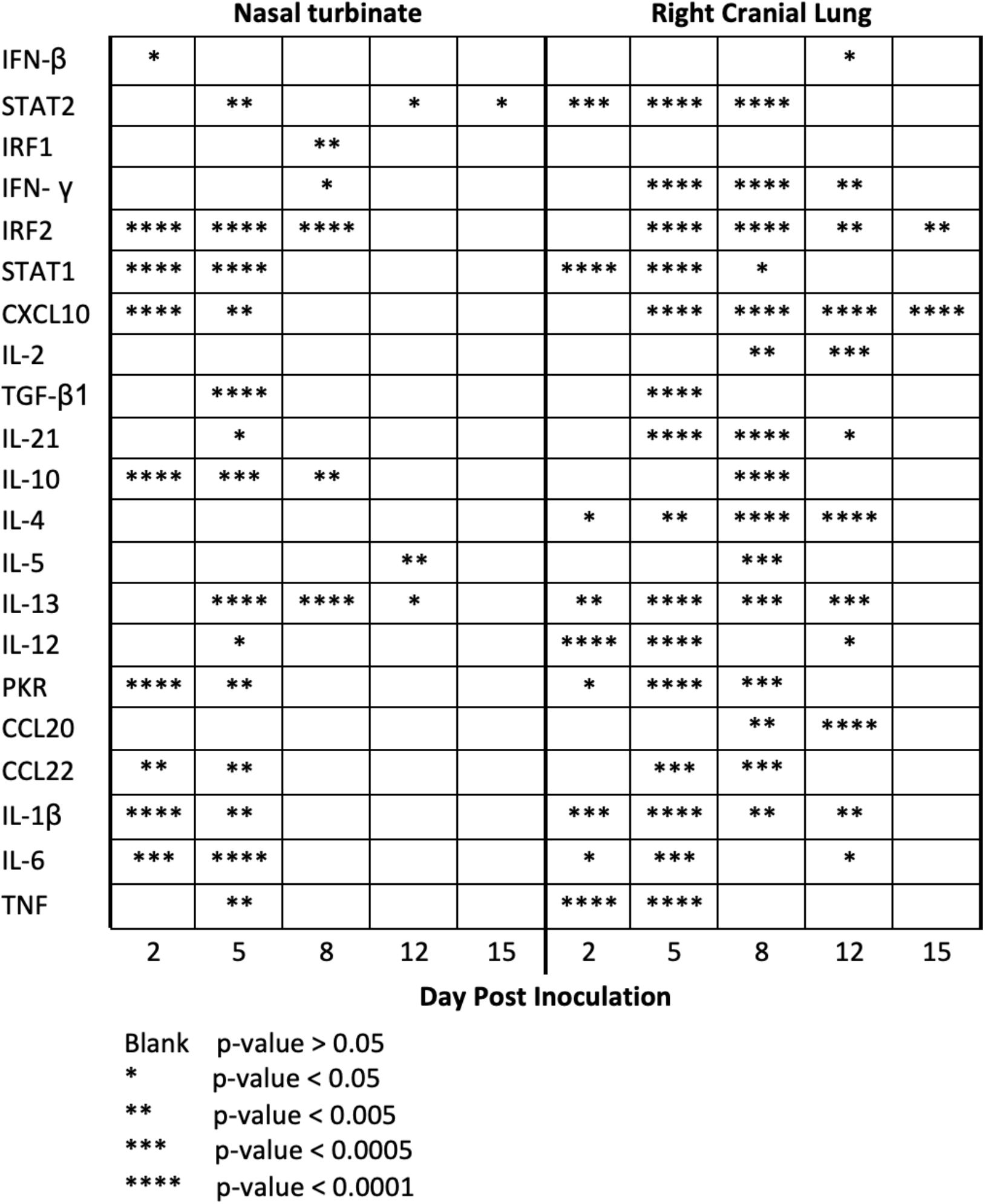
Statistical analysis of Respiratory Tissue Gene Expression Analysis. qRT-PCR was performed on RNA extracted from nasal turbinate and right cranial lung tissue from SARS-CoV-2 inoculated hamsters. Fold-change was calculated via *ΔΔ*Ct against baseline (Day 0) with BACT as the housekeeping gene **(Figure 5).** ANOVA was used to calculate statistical significance by comparing hamsters on the days post inoculation to baseline (day 0). No asterisk indicates a p-value>0.05; *indicates a p-value < 0.05; **indicates a p-value <0.005; *** indicates a p-value <0.0005; **** indicates a p-value <0.0001. N is 3 for all timepoints.

**Table S2.**
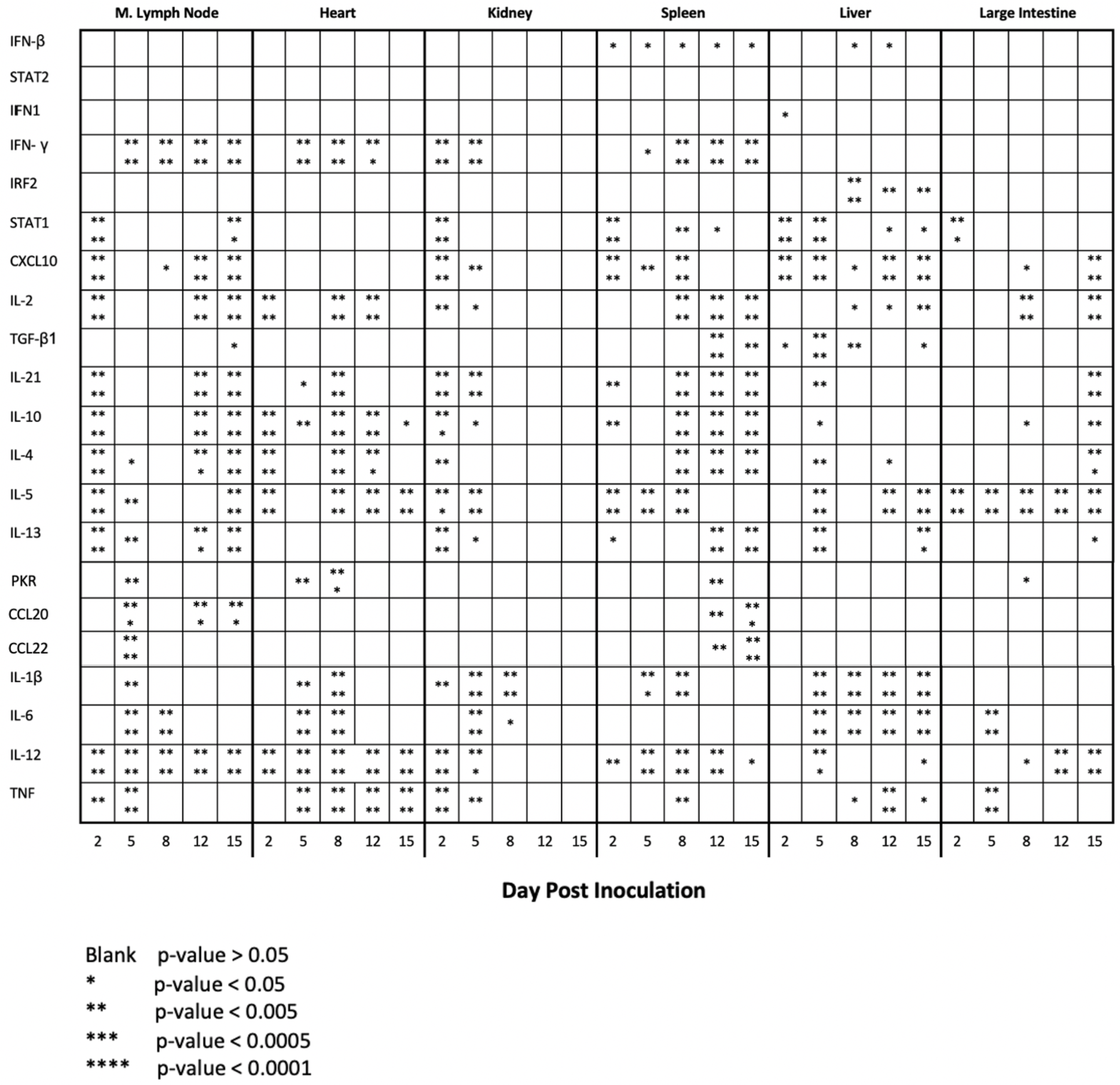
Statistical analysis of Extrapulmonary Tissue Gene Expression Analysis. qRT-PCR was performed on RNA extracted from mediastinal (M) lymph node, heart, kidney, spleen, liver, and large intestine from SARS-CoV-2 inoculated hamsters. Fold-change was calculated via *ΔΔ*Ct against baseline (Day 0) with BACT as the housekeeping gene **(Figure 6).** ANOVA was used to calculate statistical significance by comparing hamsters on the days post inoculation to baseline (day 0). No asterisk indicates a p-value>0.05; *indicates a p-value < 0.05; **indicates a p-value <0.005; *** indicates a p-value <0.0005; **** indicates a p-value <0.0001. N is 3 for all timepoints.

## Supplementary Figure Legends

**Figure S1.**
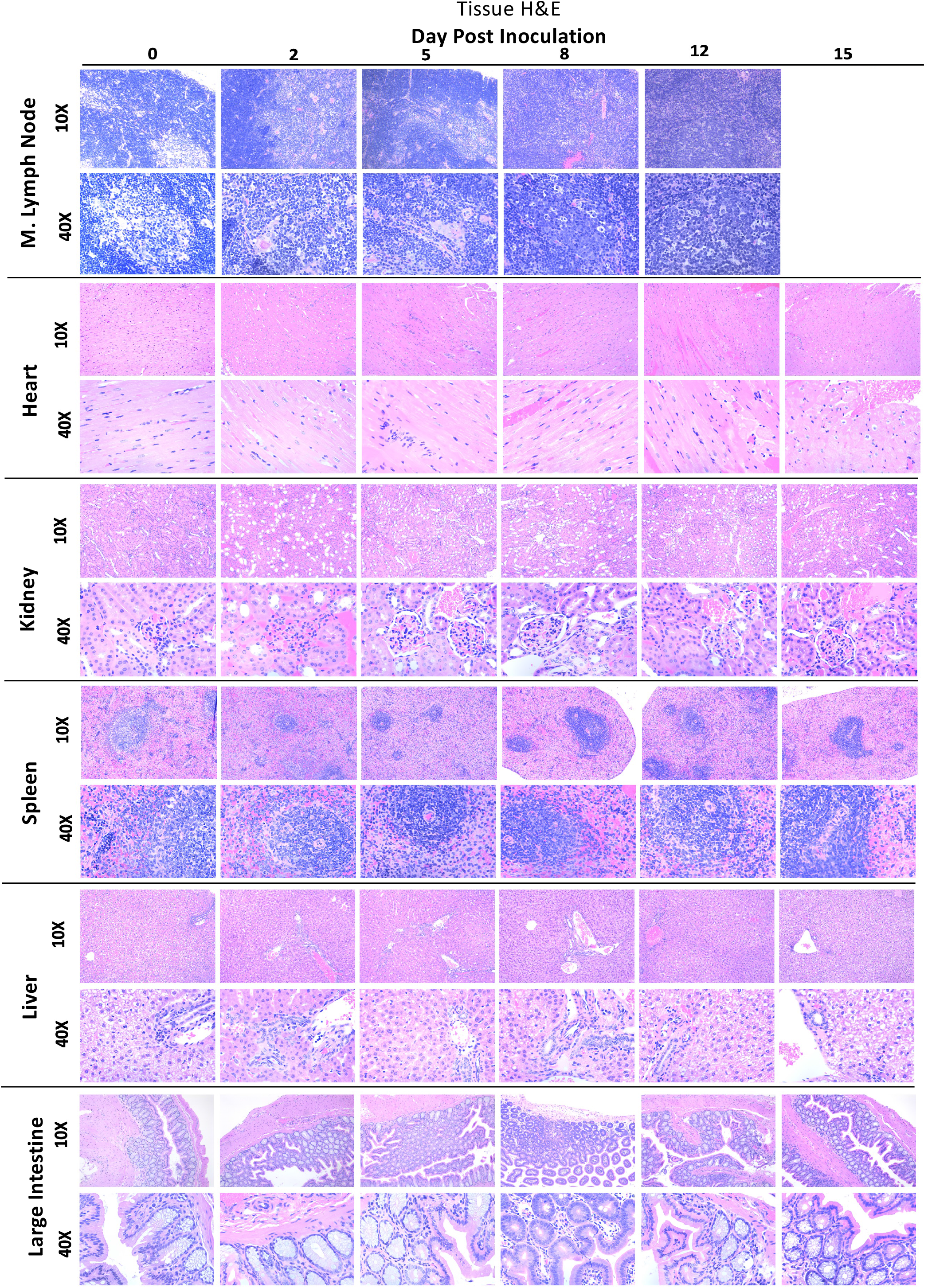
Histological analysis of tissues post SARS-CoV-2 infection demonstrate potential damage in extrapulmonary organs. The mediastinal (M) lymph node, heart, kidney, spleen, liver and large intestine were collected from each animal at necropsy and fixed in 10% formalin prior to paraffin block embedding. All tissues were analyzed by H&E. Stained tissues were visualized and imaged using the Leica DMI100 Brightfield microscope and DMC5400 20 MP color CMOS camera. Images were captured and 10X and 40X. Images shown are representative of 3 animals per group, per timepoint.

**Figure S2.**
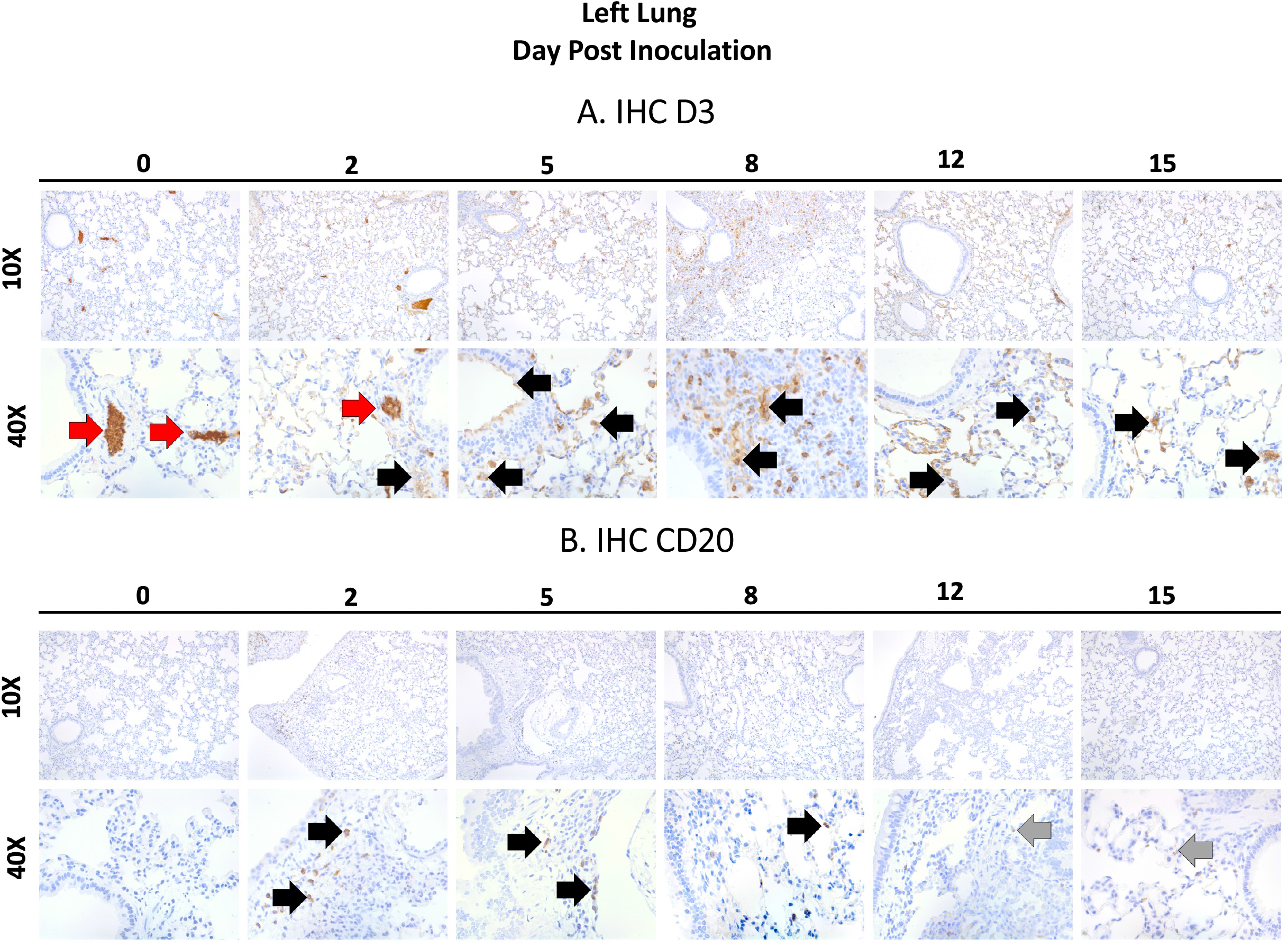
Immunohistochemical staining reveals a high infiltration of CD3+ T cells and a lower level of CD20+ B cells in the lungs of SARS-CoV-2 inoculated hamsters. The left lung was collected from each animal at necropsy and perfused with 10% formalin prior to paraffin block embedding. The lung was stained for the presence of CD3 as a marker of T cells (A) or CD20 as a marker of B cells (B). Red arrows indicate high amounts of antigen staining, black arrows indicate intermediate amounts of antigen staining, grey arrows indicate low amount of antigen staining by IHC using target specific antibodies. Stained tissues were visualized and imaged using the Leica DMI100 Brightfield microscope and DMC5400 20 MP color CMOS camera. Images were captured and 10X and 40X. Images shown are representative of 3 animals per group, per timepoint.

**Figure S3.**
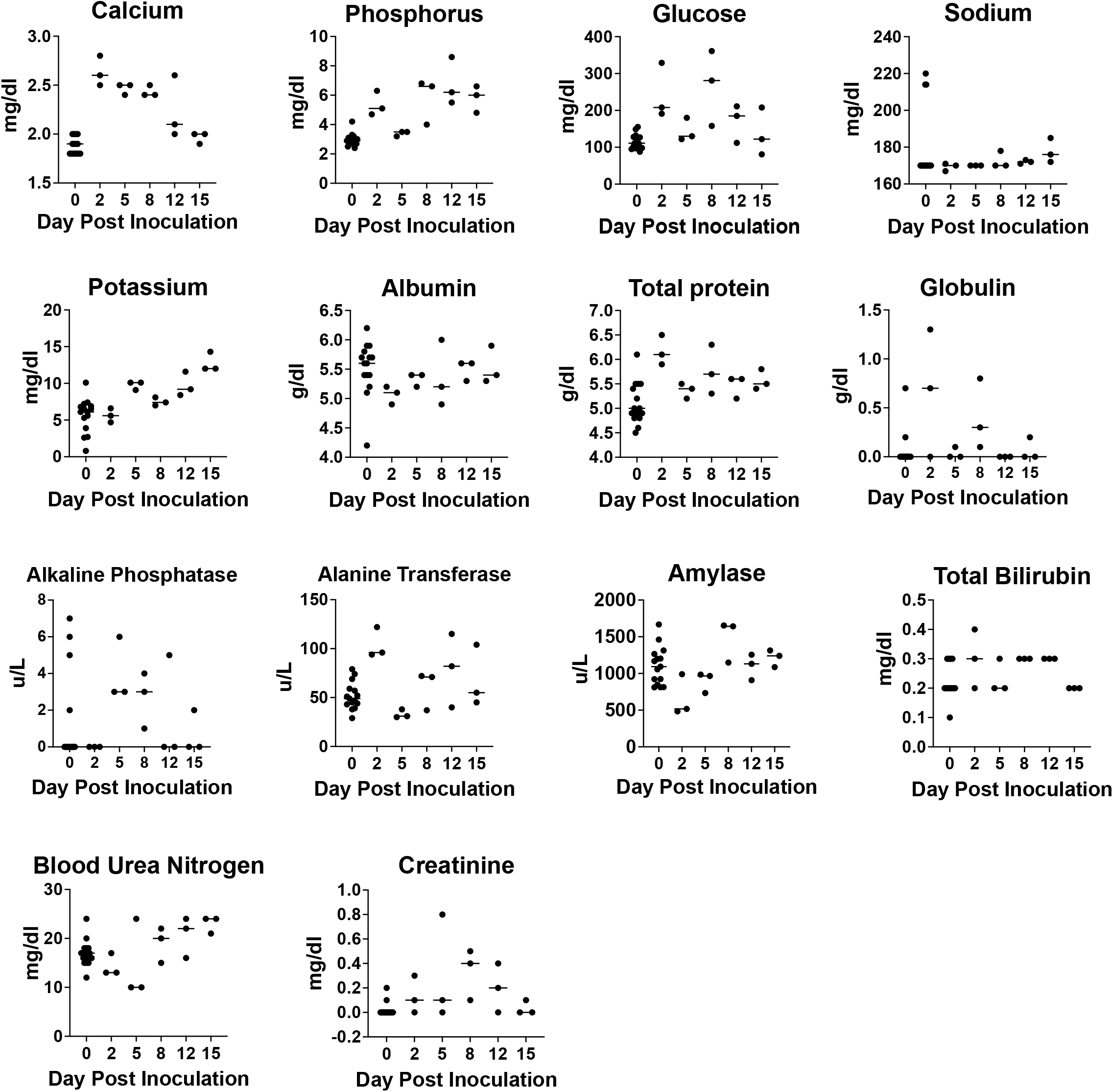
Blood plasma chemistry analysis of analyte regulation suggests potential extrapulmonary organ disruption. Plasma was separated from blood collected throughout the time course. Plasma was then assessed for 14 molecules (calcium, phosphorous, glucose, sodium, potassium, albumin, total protein, globulin, alkaline phosphatase, alanine transferase, amylase, total bilirubin, blood urea nitrogen and creatinine). Data points represent individual animals and line represents the average on a given day.

## Notes

### Competing Interest Statement

The authors have declared no competing interest.

